# Single-cell analysis of a mutant library generated using CRISPR-guided deaminase

**DOI:** 10.1101/610725

**Authors:** Soyeong Jun, Hyeonseob Lim, Honggu Chun, Ji Hyun Lee, Duhee Bang

## Abstract

CRISPR-based screening methods using single-cell RNA sequencing (scRNA-seq) technology enable comprehensive profiling of gene perturbations from knock-out mutations. However, evaluating substitution mutations using scRNA-seq is currently limited. We combined CRISPR RNA-guided deaminase and scRNA-seq technology to develop a platform for introducing mutations in multiple genes and assessing the mutation-associated signatures. Using this platform, we generated a library consisting of 420 sgRNAs, performed sgRNA tracking analysis, and assessed the effect size of the response to vemurafenib in the melanoma cell line, which has been well-studied via knockout-based drop-out screens. However, a substitution mutation library screen has not been applied and transcriptional information for mechanisms of action was not assessed. Our platform permits discrimination of several candidate mutations that function differently from other mutations by integrating sgRNA candidates and gene expression readout. We anticipate that our platform will enable high-throughput analyses of the mechanisms related to a variety of biological events.

---

Innovative biological research approaches based on the CRISPR/Cas9 system have been developed to facilitate investigations of the functional effects of genetic mutations(*1–4*). Although the functions of thousands of variants have been determined using functional screening approaches based on gene knock-out and activation, screening other genetic features, such as single nucleotide variants (SNVs) and structural variations, to examine related phenotypes remains challenging because it is difficult to introduce and analyze these genetic features for pooled screens. Among these genetic features, SNVs are the most frequently observed, and they are associated with many diseases in humans(*5*). For example, mutations related to drug resistance and cancer susceptibility significantly impact both the prognosis of therapy and disease progression. In interrogating the function of SNVs, oligo-mediated saturation editing based on homology-directed repair enables base-resolution mutagenesis(*6*), but this approach is limited to a single locus due to the difficulty of the preparation and size restriction of the oligos composed of various types of SNVs. An alternative approach based on a CRISPR RNA-guided deaminase(*7*) permits the specific and efficient mutagenesis of multiple C to T conversions in the guide RNA binding region without the need for a DNA template. Even though a small number of unintended mutations, such as indels and multiple nucleotide variants (MNV), could be introduced simultaneously, disease-related missense mutations could be generated and analyzed using this technology. However, to date, this approach has only been used for multigene knock-out(*8*) and multiple point mutations within a single gene(*9*). Introducing and screening multiple point mutations in multiple genes remains challenging due to the difficulty of analyzing multiple introduced mutations, because all mutations should be analyzed individually by amplification and deep sequencing for each locus.

As the methods described above are based on population analyses and thus dependent on the number of clones, data regarding only significantly dominant clones are obtained when using statistical algorithms(*10*). However, recently developed techniques such as CRISP-seq(*11*), Perturb-seq(*12*) and CROP-seq(*13*), which leverage single-cell RNA-seq, enable analysis of the various phenotypes and perturbations within each cell by integrating both the gene expression readout and CRISPR-based perturbations. For the perturbation read out, CROP-seq utilizes a vector that adds a poly(A) tail to the sgRNA transcript, whereas Perturb-seq and CRISP-seq generate a “perturbation barcode” linked to the sgRNA. The use of these methods has thus far been limited to investigations of perturbations associated with knock-out mutations and transcriptional regulations(*14*) generated using a CRISPR library at the single-cell level, as it remains challenging to investigate perturbations for point mutations. However, a recent report demonstrated a spectrum of subclonal point mutations in the same tumor has implications for precision medicine and immune therapies(*15*). This report emphasizes the need for an atlas of transcription profiles associated with SNVs.

To investigate the myriad of SNVs that affect biological function, a high-throughput, pooled screening method for substitution mutations which enables the analysis of generated mutations by tracking sgRNAs is required. Here, we demonstrate a novel method combining CRISPR RNA-guided deaminase and CROP-Seq technology that enables the introduction of SNVs in multiple genes and screening of the impact on function in addition to analyses of perturbations in single cells (Fig. 1A). To highlight the utility of our novel platform, we generated substitution mutations in each exon of three human genes (*MAP2K1*, *KRAS*, *NRAS*) associated with resistance to vemurafenib(*16*), which is a cancer drug targeting the *BRAF* V600E mutation in melanoma patients. We then screened for point mutations that conferred resistance to vemurafenib, analyzed the perturbations in individual resistant clones, and assessed gene expression signatures for individual clones. Finally, we selected final candidate sgRNAs, and confirmed mutation sequence by amplifying candidate regions of genome, which were extracted from edited pool of cells.

**Fig. 1.**
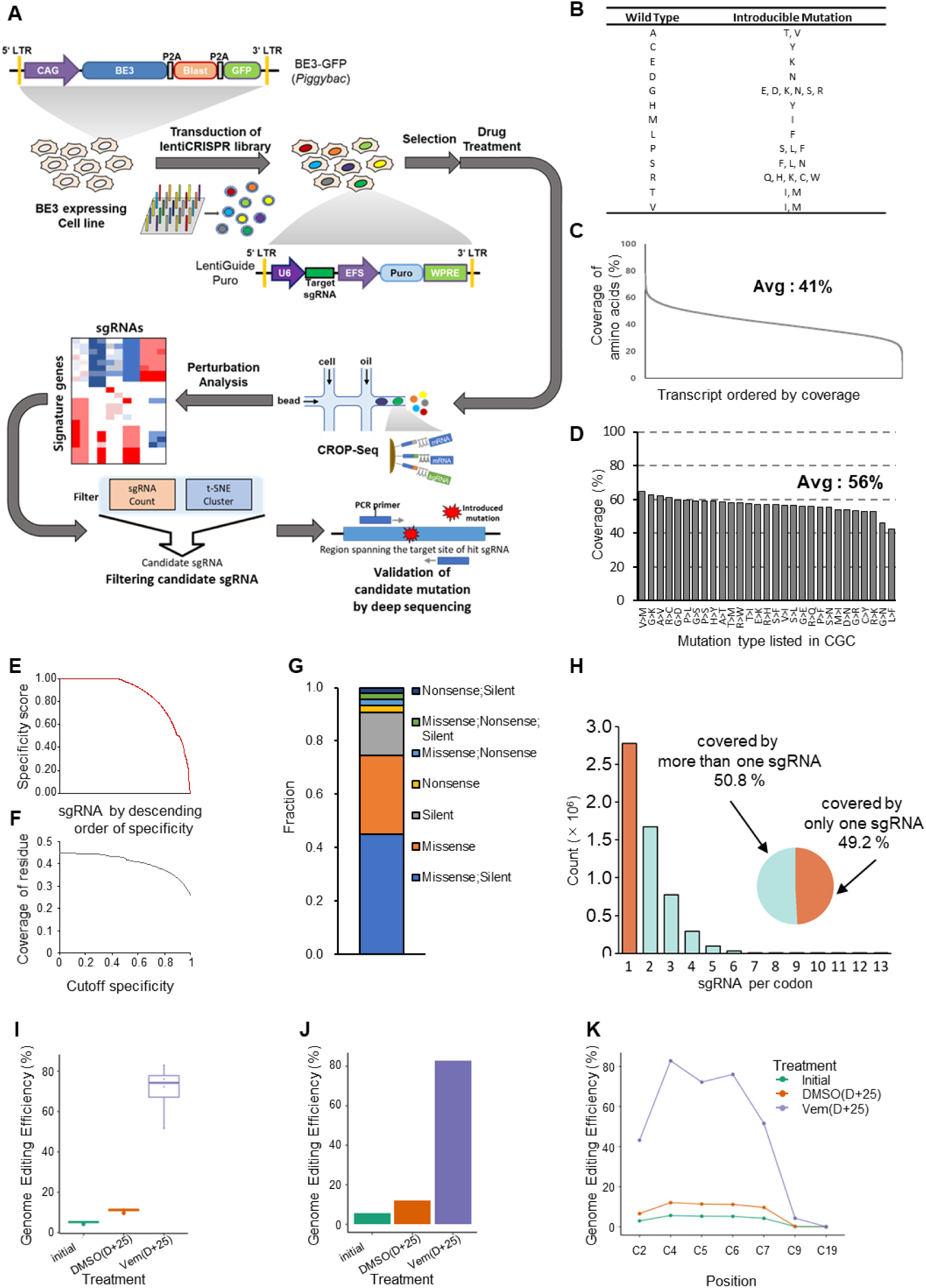
Workflow of our platform, mutations introducible using our system and demonstration of method feasibility. (**A**) Schematic flow of our platform. (**B**) Types of introducible mutations. Thirteen of the canonical amino acids can be converted to another amino acids. (**C**) On average, 41% of protein residues per gene isoform were covered by targeting “GG” and “AG” PAMs. (**D**) Introducible disease-related mutations listed in the CGC. (**E**) Specificity score for 10,038 sgRNAs designed from 20 randomly selected genes and (**F**) average coverage of residue for all possible isoforms depending on an applied cutoff of specificity score binned by intervals of 0.01. (**G**) Fraction of each introducible type of mutation. (**H**) Number and distribution of sgRNAs, indicating that more than 50% of codons can be covered by more than one sgRNA. (**I**) Targeted cytosine to thymine substitution efficiency in the “activity window” of the base editor. Solid line represents the median, with the upper and lower limits of each box corresponding to the 75% and 25% quantiles of the data. (**J**) Allele frequency of the E203K mutation before and 25 days after drug treatment. (**K**) Efficiency of cytosine substitution according to position in the protospacer of the sgRNA.

Using this platform, we could classify resistant clones into two sub-types according to drug response. Moreover, we were able to identify sgRNAs more likely to be candidates by combined analysis of candidate sgRNAs based on trackable genomic integrated sequences with transcriptome data. We anticipate that our platform can be extended to characterize individual responses related to biological stimulation in heterogeneous cells harboring single substitution mutations.

## RESULTS

### Overview of the system

To facilitate the generation and tracking of point mutations, lentivirus- and *piggyBac*-based delivery systems are needed for integration of single-copy sgRNAs in each cell and stable expression of the CRISPR RNA-guided deaminase, respectively. Because complete genes are too long for delivery by lentivirus or transposase in our system, we designed the system to utilize two vectors (Fig. 1A): lentiviral and *piggyBac* vectors encoding the sgRNA and CRISPR RNA-guided deaminase, respectively. For sgRNAs, we adopted the CROPseq-Guide-Puro plasmid(*13*) to enable simultaneous capture of the sgRNA with other mRNAs during the scRNA-seq preparation procedure and puromycin selection of sgRNA-bearing cells. For the CRISPR RNA-guided deaminase, we adopted the BE3 sequence–encoding Cas9 nickase fused with APOBEC and UGI(*4*). BE3 introduces ∼15-75% of C to T conversions with less than 1% indel formation(*7*). Although the editing efficiency of BE3 is lower than Cas9(*3*) (>90%), we believe this efficiency is high enough compared to homologous recombination-based substitution mutation-generating methods(*2*). We also constructed a cassette containing BE3 and a GFP reporter and blasticidine resistance marker, which was cloned into the *piggyBac* vector to provide high “cargo-carrying” capacity.

As BE3 can be used to introduce C to T mutations in the 4^th^ to 8^th^ positions from the PAM-distal end of the protospacer, only a limited variety of mutations can be introduced using the CRISPR RNA-guided deaminase (Fig. 1B). Thirteen of the 20 canonical amino acids can be converted to another amino acid by targeted deamination. The remaining seven amino acids can be converted only by either silent or nonsense mutations.

To simulate the extensibility and coverage of the BE system, we picked out all possible sgRNAs from all gene isoforms within the human genome (**Methods**, **Data file S1**). We observed 41% of residues per gene isoform could be targeted, if “AG” PAM(*4*) is allowed (Fig. 1C**, D**). And we observed that 45% of the sgRNAs designed from 20 randomly chosen genes (*TAL1*, *POU5F1*, *MSH6*, *CRTC3*, *WWTR1*, *CARS*, *MAP2K2*, *CDH1*, *CDKN2C*, *H3F3B*, *PTEN*, *FBXW7*, *NUP98*, *ELL*, *ITK*, *SETD2*, *TP63*, *RB1*, *CASP8*, *HMGA2*) were specific (specificity score was equal to 1, **Methods**), and the coverage of residue was 26% when sgRNAs with specificity score 1 were considered (Fig. 1E**, F**). Our simulation results indicated that one sgRNA could induce multiple types of mutations depending on the number and position of the cytosine residues in the activity window. In most cases, silent and missense mutations could be produced alone or in combination with sgRNA (Fig. 1G). We also observed 50.8% of loci were covered by more than one sgRNA; therefore, more than one sgRNA could be selected from the candidates for a more robust statistical analysis (Fig. 1H).

We utilized the above-described system to screen for mutations conferring resistance to vemurafenib to demonstrate that the system is suitable for large-scale screening of SNVs. Although mutations conferring resistance to vemurafenib have been thoroughly studied using approaches such as drop-out knock-out screening(*3*), to date, there are no reports of systematic analyses of transcriptional changes induced by various SNVs. We therefore explored how such mutations affect the drug-response mechanism using our novel CRISPR RNA-guided deaminase system.

### Introduction and enrichment of resistant mutation MAP2K1 E203K

To assess the efficiency of our system, we investigated the rate of conversion of the C base targeted in a single locus. First, we introduced the *MAP2K1* p.E203K mutation (c.G607A) into human A375 melanoma cells carrying a homozygous *BRAF* V600E mutation (c.T1799A). The E203K point mutation results in the substitution of glutamic acid with lysine at codon 203 in exon 6 of *MAP2K1*, resulting in resistance to vemurafenib.

The *piggyBac* vector expressing BE3 was transfected into A375 cells to create a cell line that stably expresses BE3. Cells expressing the base editor were infected with a lentivirus expressing sgRNA, targeting codon 203 of *MAP2K1* to induce the E203K mutation. The cells were cultured for 10 days to ensure selection of cells harboring the sgRNA and to promote base editing. Then, frequency of base editing in terms of base and amino acid resolution was investigated by deep sequencing after 25 days of treatment with either vemurafenib or vehicle (dimethyl sulfoxide [DMSO]). At 10 days after puromycin selection, the substitution efficiency ranged from 4.24–5.65% in the “activity window” with a small indel frequency of 1.4% (Fig. 1I). After treatment with vemurafenib for 25 days, the substitution efficiency increased significantly, to 41.11–77.17%, whereas DMSO vehicle treatment yielded an efficiency of only 9.67–12.08%. These data suggest that continuous engineering occurred during culture and that the mutation confers resistance to vemurafenib. Therefore, cells harboring the E203K mutation survived over wild-type cells in the presence of vemurafenib, whereas DMSO treatment did not promote enrichment of the mutant clones as much as vemurafenib. Consistent with the base resolution analysis, codon-based analysis indicated that 77.19% of the viable cells were E203K clones (Fig. 1J). The same analysis of DMSO-treated cells revealed that 12.06% of the viable cells were E203K clones. These results confirmed that significant enrichment of the mutation occurred with vemurafenib treatment. As shown in a previous study(*7*), conversion of C residues can occur at a variety of positions, although conversion occurs most frequently within the activity window (Fig. 1K, fig. S1A). Furthermore, processive deamination(*7, 17*) tends to occur in same DNA strand (fig. S1B). Collectively, the results of our singleplex experiments suggest that the mutations artificially introduced via base editing functioned well and that mutants can be specifically enriched using drugs in a manner similar to naturally acquired mutations in patients. These data also demonstrate the possibility of using this system to measure the effect sizes of various mutations in terms of drug resistance.

### Introduction and functional screening of multiple putative vemurafenib-resistance mutations using population analyses

To determine whether the use of our system can be extended to the analysis of multiple loci, we designed a sgRNA library for three genes (*MAP2K1*, *KRAS*, and *NRAS*) related to vemurafenib resistance in melanoma. We selected possible targets based on the criterion that spacers include cytosine residues in the activity window. A total of 420 sgRNAs were designed for all of the exons of *MAP2K1*, *KRAS*, *NRAS* (263, 80, and 77 sgRNAs, respectively, **Data file S2**, fig. S1C). We excluded the sgRNAs used in the previous singleplex experiments to avoid enrichment of the known E203K resistance clone in the pooled screen. The resulting sgRNA library covered 17.4% of the reported disease-related SNVs in the CGC (Fig. 2A). Among a library of sgRNAs, we independently tested three sgRNAs by viral production and transduction and compared with non-targeting sgRNA. We observed an editing efficiency of 0.40 – 7.85 % of editing efficiency in the ‘activity window’ and very small portion of indels (< 0.09%) for these three sgRNAs (**Methods**, fig. S1D–J). Also, it appears that there are no significant false-positive effects on resistance (**Methods**, fig. S1K, L). The designed library was then synthesized using a microarray and cloned into the CROP-guide-puro plasmid. A375 cells stably expressing BE3 were transduced in two independent replicates with the library and selected for 14 days using puromycin to ensure that most of the cells expressed sgRNA and were base edited. Before drug selection, we determined the C to T conversion rate in sgRNA library transduced cells. The conversion rate was not as high as a singleplex experiment because total editing efficiency was divided by 420 loci. However, the conversion rate was significantly higher than the sequencing error observed in non-targeting sgRNA-transduced cells (fig. S2A–C, **Methods**). Then, the cells were treated with either vehicle (DMSO) or vemurafenib for 28 days. Cells were sampled on the day treatment was started and 7, 14, 21, and 28 days after treatment (hereafter designated initial, D+7, D+14, D+21, and D+28). Before examining the transcriptome of single cells, the population of sgRNAs integrated into the genome was analyzed to determine which sgRNA was responsible for conferring resistance.

**Fig. 2.**
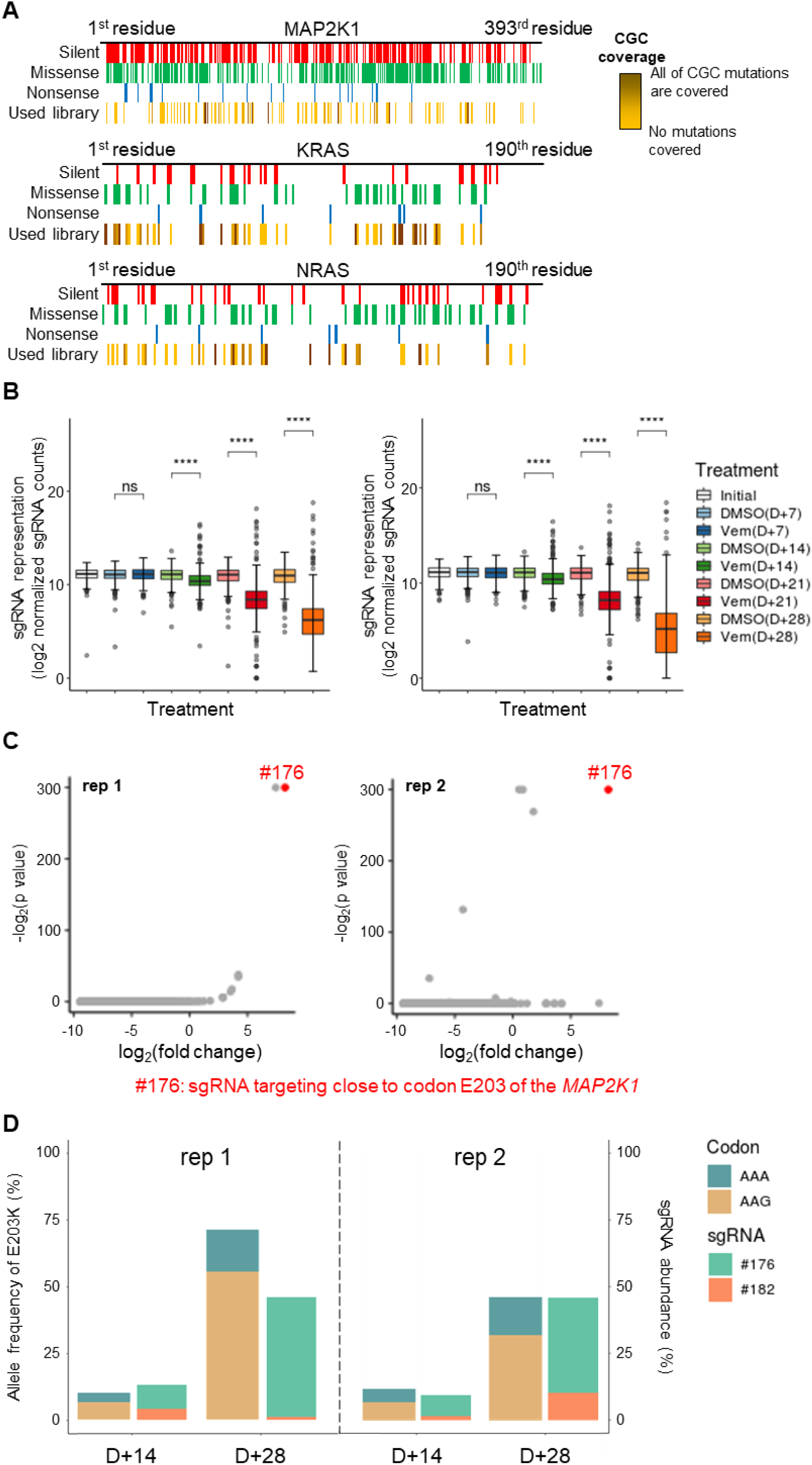
Introduction and functional screening of multiple mutations using population analysis. (**A**) Introducible disease-related SNVs in a sgRNA library targeting *MAP2K1*, *KRAS*, and *NRAS*. Introducible silent, missense, nonsense, and CGC mutations are shown in red, green, blue, and yellow, respectively, according to residue position in the protein. Coverage of CGC mutations is indicated by a separate color (maximum coverage = 100%, indicated by dark yellow). (**B**) Box plot showing the distribution of reads from individual sgRNAs by time (initial, D+7, D+14, D+21, D+28) and condition (DMSO, vemurafenib [Vem]) in replicate 1 (left) and replicate 2 (right). All p-values determined by two-sided Wilcoxon rank-sum test, ****p < 0.0001. ns: not significant (p > 0.05). The center line corresponds to the median; the edges of the box correspond to the first and third quartiles; the whiskers represent 1.5× the interquartile range; and the black dots indicate outliers. (**C**) Scatter plot showing enrichment of sgRNAs with their corresponding p-values. #176 sgRNAs targeting close to E203 of MAP2K1 are shown as red dots. P values less than 10^-300^ were arbitrarily fixed to 10^-300^ for easy comparison of the sgRNAs. (**D**) Boxplot showing the allele frequency of the E203K clone and abundance of sgRNA-introduced E203K mutation in *MAP2K1* from experiment replicates 1 and 2.

First, the distribution of sgRNAs for each condition was investigated. The sgRNA representation decreased over time, meaning that some sgRNAs were enriched over time (Fig. 2B). An analysis using MAGeCK(*10*) indicated that sgRNA #176, which targets close to codon E203 in the *MAP2K1* gene, was enriched in both independent screens (Fig. 2C). It should be noted that sgRNAs targeting close to codon E203 (#176, #182) accounted for ∼45% of all sgRNAs, and E203K mutant cells constituted over 70% of the total cell population in replicate 1, suggesting that the abundance of sgRNAs close to E203 is reflective of the frequency of E203K mutant cells (fig. S2D, Fig. 2D). Although we excluded the highly active sgRNA generating E203K, the data suggested that other sgRNAs in which the protospacer included the E203 residue in the extra position of the activity window robustly generated the E203K mutation. In sgRNA #176, the E203 residue is in the 13^th^ to 15^th^ position from the PAM-distal end of the protospacer, such that the E203K mutation is predominant compared with the mutation in the editing window due to the TC motif preferred by the APOBEC1 enzyme in BE3(*7*). The E203K mutation was also detected in replicate 2, with an allele frequency of ∼48% (Fig. 2D, fig. S2E). These results demonstrate that generating a library comprised of multiple mutants is possible with our novel system and that appropriate targets can be selected using our platform. However, further analyses using transcriptional data are warranted in order to identify other putative sgRNAs and elucidate the mechanism of enrichment.

### Identifying cell subpopulations across different treatment periods using scRNA-seq

The scRNA-seq approach was employed to explore properties related to drug responses in individual cells. To assess the transcriptional changes occurring in each mutant, cells from each experimental condition (before treatment: initial; 14 days after DMSO or vemurafenib treatment: DMSO(D+14) or Vem(D+14); 28 days after DMSO or vemurafenib treatment: DMSO(D+28) or Vem(D+28)) were individually harvested and subjected to Drop-seq(*18*). The mRNAs and sgRNAs in each cell were captured and converted into a cDNA library for NGS. On average, we sequenced 82 million reads per sample (Table S1). After filtering cells in replicates 1 and 2, 5707 and 7511 cells with more than 500 genes were identified, respectively, and 57.4 and 55.2% of the cells were assigned as sgRNAs, respectively. These results were comparable to data from a previous report(*13*).

We first compared the abundance of each sgRNA to the abundance determined from genomic DNA obtained from bulk cells. High correlations were observed between the sgRNA and genomic DNA data (Fig. 3A). In particular, enrichment of sgRNAs introducing the E203K mutation was observed in drug-treated samples from both analyses, indicating that the transcriptome data were reliable for analyzing the perturbations associated with the assigned sgRNAs. To confirm transcriptome concordance, we investigated whether the Euclidean distance between cells with the same sgRNAs in a different replicate is smaller than the distance to other sgRNAs. We observed that the distance to different replicate is smaller than distance to different sgRNAs (fig. S2F). Similarly, the distance between sgRNAs with identical target strands and overlapping protospacer regions was smaller than different sgRNAs targeting different regions (fig. S2G).

**Fig. 3.**
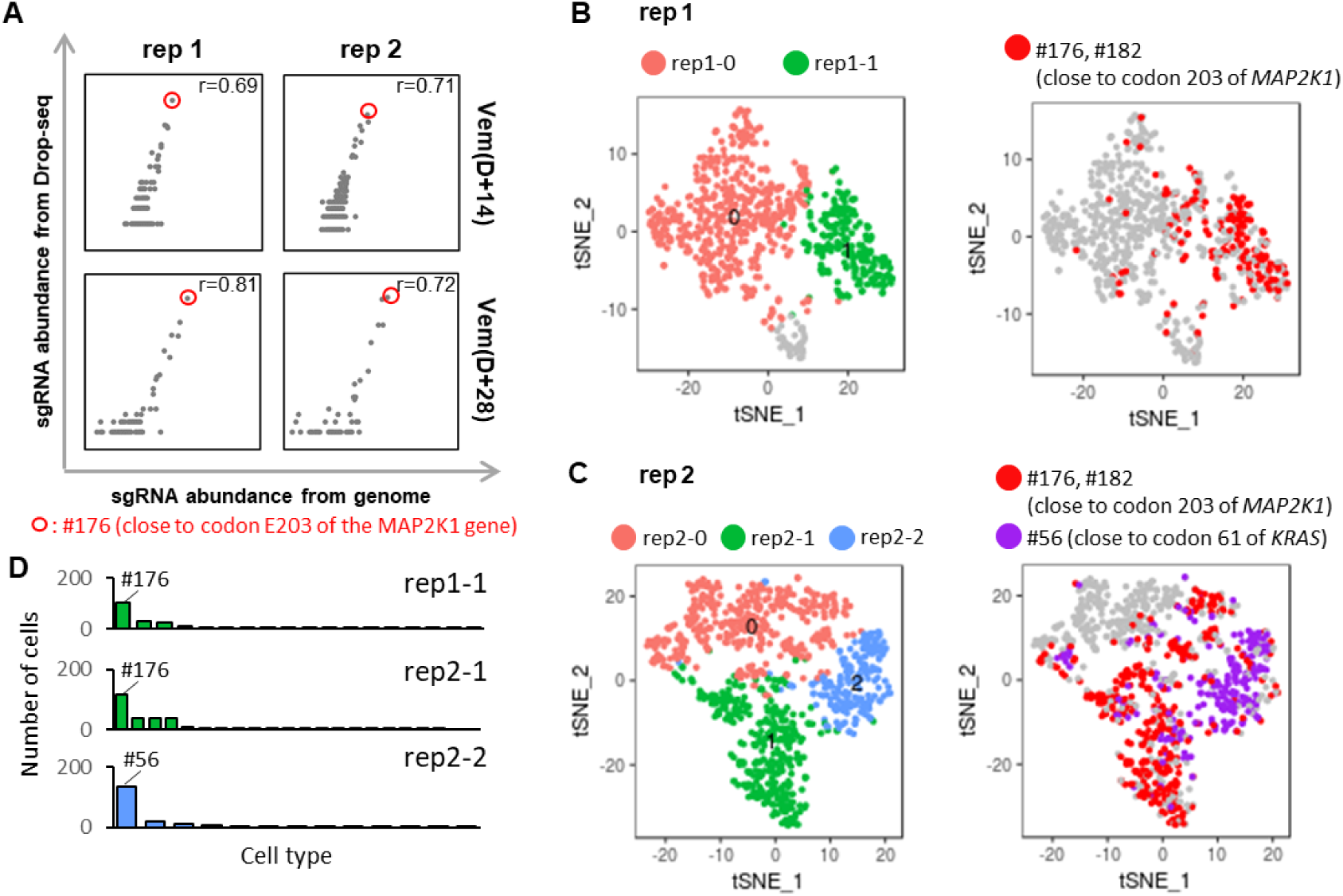
Analysis of abundance and clusters in scRNA-seq data. (**A**) Concordance of sgRNA abundance from Drop-seq data and genomic data for each replication and condition. The #176 sgRNA, which potentially introduces a known mutation (E203K), is marked by a red circle, and the Pearson correlation coefficient is shown in the upper right corner of the plot. (**B**) t-SNE visualization of D+28 cells (left) and distribution of #176 sgRNA-containing cells (right) in replicate 1. (**C**) t-SNE visualization of D+28 cells (left), distribution of #176 and #56 sgRNA-containing cells (right) in replicate 2. (**D**) Distribution of sgRNAs for rep1-1, rep2-1, and rep2-2.

We also estimated MOI and detection rate using the equations in Dixit et al.(*12*) to determine whether sgRNA detection was reliable (**Methods**). We observed a 66% detection rate with an inferred MOI of 0.83, and a 41% detection rate with an inferred MOI of 0.98, for initial cells of replicate 1 and 2, respectively (figs. S2H, I). We expect that low detection rate was due to the low complexity of the cells’ profiled (cells with more than 500 genes detected were considered, and the median number of detected genes for these cells was 894 and 961.5 for each replicate, respectively). Despite the low detection rate, our observations were comparable with those of Dixit et al.

Next, we performed a principal component analysis (PCA) and trajectory analysis to determine whether there is a characteristic cluster pattern according to period and treatment. The transcriptome of resistant cells at D+28 and D+14 was distinct from that of naïve cells in each replicate (fig. S3A). However, no common pattern was observed between replicates (fig. S3B, C). Also, an attempt was made to identify significant relationships between period and condition by linear regression with the ordinary least square (OLS) model (**Data files S3, S4**). However, we could not observe a significant relationship (fig. S3D) even though many genes were predicted to show linearity between treatments (DMSO vs. vemurafenib). Most of genes varied depending on condition rather than sgRNA-specific responses. Therefore, we focused on D+28 cells to assess the mutational effects of sgRNAs in more detail.

D+28 cells were visualized using t-SNE and grouped by unbiased clustering (Fig. 3B**, C**). We hypothesized that the transcriptome pattern of cells that acquired resistance due to the E203K mutation would differ significantly from that of natural survivors. Thus, we expected that the cluster including the #176 sgRNA introducing E203K would be separated from other clusters. In the t-SNE visualization, we observed two and three visually distinct clusters based on a neighborhood graph defined by Euclidean distances in PC spaces from replicates 1 and 2, respectively. One cluster in both replicate screens (rep1-1, rep2-1) consisted primarily of #176 sgRNA, which targets close to E203 in *MAP2K1* (Fig. 3D, fig. S4A). In replicate 2, an additional cluster composed primarily of #56 sgRNA targeting close to Q61 of *KRAS* (rep2-2) was identified (Fig. 3D, fig. S4B). We hypothesized that these clusters consisted of cells that had acquired resistance and therefore investigated them further to elucidate the resistance mechanism in more detail.

### Investigation of cluster of resistance-acquired cells

We investigated whether transcriptional changes in the clusters composed primarily of #176 sgRNA targeting close to E203 in MAP2K1 were related to vemurafenib resistance by identifying marker genes. Among candidate up-regulated marker genes identified using the Wilcoxon rank sum test, genes were not differentially expressed in DMSO-treated cells except for four genes (**Data file S5,** Table S2, fig. S4C-F). We expect that the marker gene expression was mostly sensitive to the treatment condition. The top 10 genes with lowest p-values were selected as “signature genes” (**Data file S6,** Fig. 4A, fig. S4G). Gene ontology (GO) and pathway enrichment analyses(*19, 20*) of the signature genes showed that the clusters were enriched with gene sets related to immune responses such as antigen processing and presentation of peptides via MHC class II (Fig. 4B). This observation was similar to the results of a previous report(*21*). *CD74*, *HLA-DRA*, *SLC26A2*, *HLA-DRB1*, *FOS*, and *HLA-DPA1* were commonly identified as signature genes in the clusters from both replicates (Fig. 4C). When we extended the criterion for marker genes to include all listed genes with a p-value <0.05, a total of 66 up-regulated genes and 163 down-regulated genes were identified as common (Fig. 4C, fig. S5, **Data file S6**). This result indicates that the members of these clusters are similar and that their perturbations are reproducible. We hypothesize that most members of these clusters are perturbed by the E203K mutation and associated with immune responses.

**Fig. 4.**
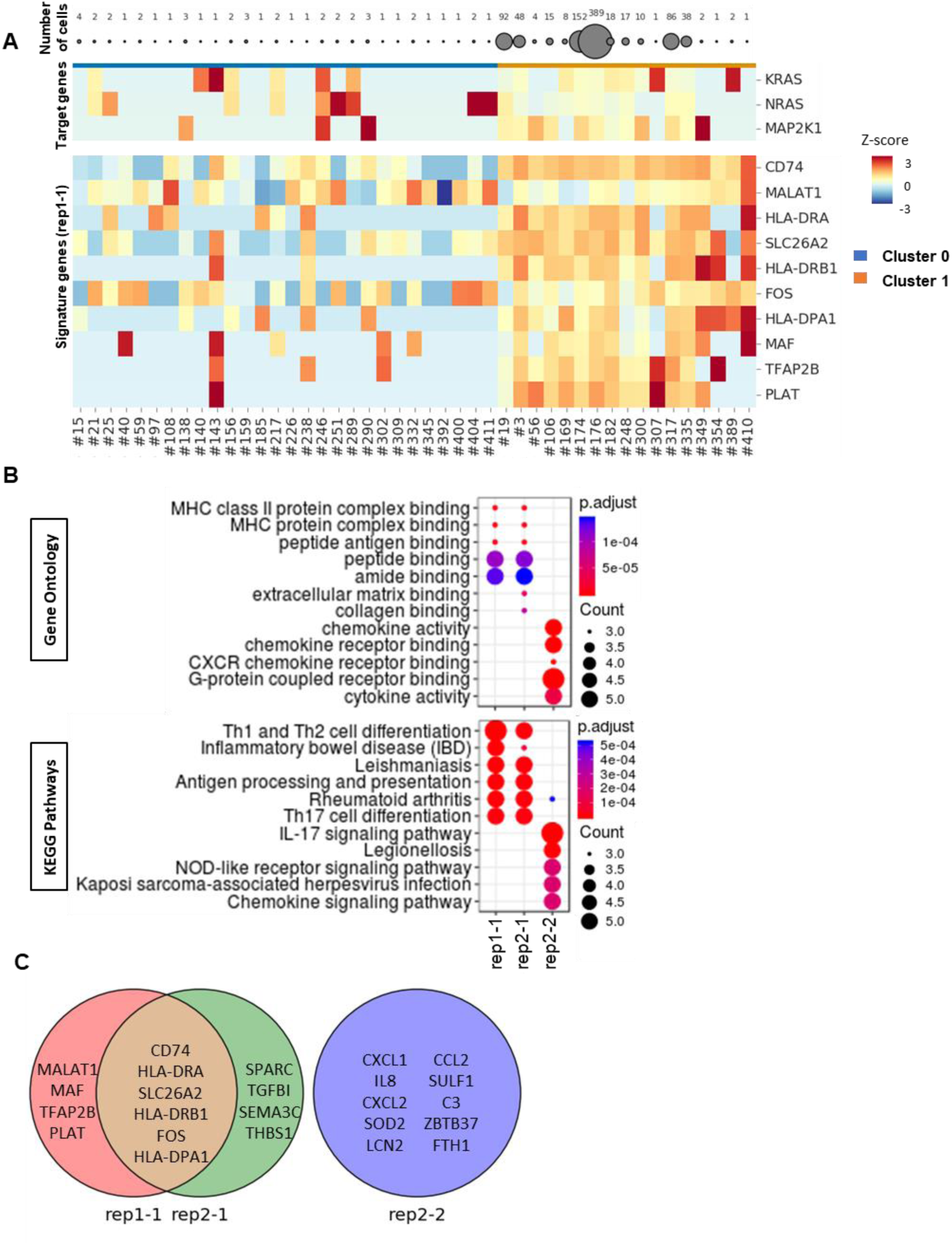
Transcription profiles of signature genes from clusters composed primarily of #176 sgRNA. (**A**) Average expression of signature genes according to individual sgRNA. Each row represents one of the signature genes, and each column represents sgRNA-bearing cells. Vem(D+28) cells in other clusters and Vem(D+28) cells in rep1-1 are arranged from left to right. Color scale indicates standard deviation of gene expression from the mean expression value, with red indicating high expression and blue indicating low expression level. (**B**) Gene ontology and pathway enrichment analyses of marker genes. Partial list of the significantly enriched GO Molecular Function (top) and KEGG Pathways (bottom). (**C**) Venn diagram of signature genes from rep1-1, rep2-1, and rep2-2.

The rep2-2 cluster was composed primarily of #56 sgRNA, which targets close to Q61 of *KRAS*, whereas representation of #176 sgRNA was low (Fig. 3C). Up-regulated marker genes (**Data file S6**) did not overlap with differentially expressed genes in DMSO-treated cells (**Data file S5,** Table S2). The top 10 genes with the lowest p-values were selected as signature genes. Ontology and pathway enrichment analyses were then performed. The signature genes of this cluster (i.e., rep2-2) differed completely from those of the rep1-1 and rep2-1 clusters (Fig. 4C**, Data file S6**). Although their expression was not as significant compared with rep1-1 and rep2-1 (fig. S4G), the signature genes were partially enriched in GO terms and pathways associated with chemokine signaling (Fig 4B). It has been reported that activated CXC chemokine receptor (CXCR) signaling in melanoma cells contributes to vemurafenib resistance(*22, 23*). Our results indicate that the transcriptional changes in the rep2-2 cluster are distinct from those of the rep1-1 and rep2-1 clusters. We hypothesize that the cells in the rep2-2 cluster composed primarily of #56 sgRNA survive via a mechanism different from that by which cells in the rep1-1 and rep2-1 clusters mainly composed of #176 sgRNA survive.

We next investigated whether other sgRNAs are common to rep1-1, rep2-1, and rep2-2. We identified nine, eight, and five sgRNAs with more than two cells in rep1-1, rep2-1, and rep2-2, respectively (Table S3). Of these sgRNAs, two (#176, #182) that could introduce E203K in *MAP2K1* were common in these clusters, and the #56, #126, and #217 sgRNAs account partially or almost completely for the rep2-1 and rep2-2 clusters (fig. S6A). Other sgRNAs were minor components (1.4% on average). Mutations that can be introduced by these sgRNAs (#56, #126, and #217) are thus potential candidates for conferring resistance to vemurafenib. We next examined whether these sgRNAs introduce cognate mutations (Table S4).

### Validation of genomic loci targeted by candidate sgRNAs via deep sequencing

We performed deep sequencing of genomic loci targeted by the candidate sgRNAs to verify the introduced mutations. Genomic loci of the #56, #126, and #217 sgRNAs were sequenced. An indel mutation (7.7%) was introduced in the target region of #56 sgRNA (fig. S6B). Because we used BE3, which employs Cas9 nickase, the indel mutation could be introduced into a small proportion of cells. There were two main types of in-frame indel mutations introduced (c.171insCTGTTGGATATTCTCGAC and c.176delCAG), and no substitutions were observed (fig. S6B, C). Enrichment in the indel mutations was observed compared to control cells treated with DMSO (0.03%). We also observed that indel introduction via lentiCRISPR system with the same #56 sgRNA conferred vemurafenib resistance. Although the total frequency of indels decreased, in-frame insertions were enriched (3.21-fold average, standard error of mean = 0.895 in triplicate experiments) during vemurafenib treatment compared to DMSO treatment (fig. S6E-I). We speculated that cells with the #56 sgRNA that introduced indel mutations next to the *KRAS* Q61-encoding sequence conferred resistance to vemurafenib. In contrast, deep sequencing of the target regions of the #126 and #217 sgRNAs did not confirm introduction of cognate mutations (fig. S6J, K). We also observed weak editing ability for #126 and #217 sgRNAs, and edited cells were not enriched in single locus editing experiments (fig. S6L, M). We believed that other biological processes such as epigenetic reprogramming(*24*) or RNA editing via rAPOBEC1(*25*) should be investigated to explain inconsistent survival during pooled screening.

In summary, we first observed enrichment of the #176 sgRNA in sgRNA abundance analyses and considered the #56, #126, and #217 sgRNAs as additional candidates as a result of transcriptome analyses. Of these selected sgRNAs, those targeting close to the sequences encoding E203 of the *MAP2K1* gene and Q61 of the *KRAS* gene (#176, #56) generated cognate mutations and induced different transcriptional changes.

## Discussion

In the present study, we established a platform combining CRISPR RNA-guided deaminase and scRNA-seq technologies that enables the introduction of substitution mutations into multiple genes and facilitates their functional screening and measurement of perturbations in single cells. We demonstrated the introduction of substitution mutations into each exon of three genes and screened for substitution mutations conferring resistance to vemurafenib using population analyses.

The results of population analyses indicated an enrichment of the E203K substitution mutation in *MAP2K1*, consistent with a previous study. Even though the pattern of substitution mutations (c.607 G>A, c.609 G>A) is different with naturally occurring substitution (c.607G>A), variants which generate the same amino acid change (E203K) were enriched under vemurafenib selection. This result highlights that our platform could be used for screening clinically actionable SNVs. Screens were also done in a simple and cost-effective process compared to arrayed screens. This highlights that our platform has the same advantage as other pooled screening methods(*26*). In addition, by employing scRNA-seq technology, we identified the perturbation as well as the signature of SNVs at the transcriptome level. We observed activation of genes associated with immune response with the E203K mutation introduced into cells, and activation of chemokine signaling in indel-introduced cells. The indels were especially focused at the D57 – G60 residues of the KRAS gene and in-frame indels were dominant over than frame-shift indels. These mutations were not reported in previous studies, but region in which the mutations were introduced is next to the sequence encoding the Q61 residue of K-Ras, which is involved in constitutive activation of intrinsic GTPase activity(*27*). Activated Ras is known to positively regulate the expression of various chemokines and ultimately promote tumorigenesis(*28*), which is consistent with the present result demonstrating that chemokine signaling pathways and CXCR binding genes were enriched in clusters composed primarily of the #56 sgRNA.

We believe that validation procedure is very important to avoid mis-interpretation because unintended mutations could also be enriched by the selection such as the indels introduced by sgRNA #56 in the *KRAS* gene. A recent study using targeted AID reported the introduction of multiple SNVs and assessed the effect of the introduced SNVs on the mechanism of imatinib resistance(*9*). The introduced SNVs were confirmed and validated by amplification of targeted exons using genomic DNA. However, this technology was hard to expand to include multiple genes or multiple exons, because the technology requires amplification of individual regions for confirmation. Multiple PCR reactions should be performed to confirm individual loci. On the other hand, our platform utilized a lentivirus-expressing sgRNA that can be tracked by single amplification of the integrated sgRNA so that we could expand our technology to include all exons in three genes. Combining this sgRNA approach with transcriptome analysis, we could narrow the scope by filtering out marginal sgRNAs. In this way, only a small number of PCR reactions which target the selected candidates are required to confirm which mutation was introduced.

Improved engineering efficiency could provide for more-accurate assessment of the perturbations associated with individual sgRNAs. A knock-out–based study reported that approximately 70% of sgRNA-bearing cells can be edited using sgRNA(*3*). In contrast, experiments using BE3 indicated that 5–20% of sgRNA-bearing cells can be edited using sgRNAs(*7*). These data suggest the possibility of discordant transcriptomic changes resulting from use of the same sgRNA, which could create confusion in analysis. Optimization of the engineering efficiency can be achieved using BE4 or BE4max(*29*). By achieving a higher efficiency in base editing (i.e., increasing the likelihood that sgRNA-bearing cells will be engineered), perturbations in a larger number of cells can be characterized. In addition, we expect that the use of a higher throughput device such as 10x Genomics Chromium would help to more accurately assess the perturbations after filtering marginal cells.

We observed C to T conversion outside the ‘activity window’ for #176 in the multiplex experiment. This conversion took place to a similar degree as ‘C’ inside the ‘activity window’, but the cells with conversions outside were enriched during vemurafenib selection. This conversion was presumably because the ‘TC’ motif is preferentially edited by BE3(*7*). Therefore, we should consider C to T conversions outside of the ‘activity window’ when designing a screening experiment.

We envisage that our approach could be optimized further by employing an alternative to BE3 protein. Recently developed BE variants and other effector proteins(*29*) provide a broader range of targets, thus increasing the analytical coverage of diseases related to SNVs. Off-target editing due to the CRISPR or non-specific deaminase activity would also be considered. A recent report(*30*) showed BE3 induces off-target SNVs on DNA and RNA, even if the loci were not potential off-targets of the guide sequence. These off-targets could hamper interpretation of mutations related to the function of interest. Replacement of BE3 with a variant deaminase could reduce the off-target activity(*29*). In addition, we observed heterogenous populations of edited cells through processive deamination (fig. S1B, S1H–J). Although cells with specific mutations that affect protein function are expected to be enriched during screening, using another base editor(*29, 31*) would permit reduced processivity and unwanted C to T conversions in the target strand, generating a less heterogeneous population of edited cells. We believe that more comprehensive analyses of mutations present could be achieved by optimizing protein selection.

We used oligo-dT sequence–linked beads to capture polyadenylated mRNAs in the scRNA-seq experiment. By synthesizing target sequence–linked beads, targeted sequences could be captured directly without laborious PCR amplification of each targeted sequence(*32*). Alternatively, ligating target sequences to conventional beads could also be used(*33*). Another application involves coupling of methods targeting individual cells using barcoding(*34*). A target cell’s cDNA can be amplified using this method in order to analyze the transcriptome, and the method is applicable to the study of cells with known mutations and validation of cells enriched with specific sgRNAs. Furthermore, although we demonstrated the effect of SNVs introduced into exons of three genes in the present study, further exploration of other exons in genes related to cancer(*5*) is expected to accelerate the identification of cancer-driving and drug-resistance mutations. Unlike the case of knockout screens resulting in similar phenotypes independent of the target residues along the coding sequence, phenotype of substitution mutation is very diverse, dependent upon the residue and mutation type even if the targeted gene is same.

According to our simulation, our system could cover 36,211 missense mutations and 3,491 nonsense mutations among the mutations listed in the Cancer Gene Census (CGC)(*5*). This indicates that a large proportion of known cancer-related mutations could be generated and examined using our system. When considering only the possible types of missense mutations that can result from conversion of the 13 canonical amino acids, our system covered 56% of the mutations listed in the CGC (Fig. 1D). We expect our platform to be extended to the examination of substitution mutations related to a variety of biological responses.

## Methods

### *piggyBac*-BE3-GFP: Construction and establishment of BE3-expressing cells

The *piggyBac*-BE3-GFP plasmid was constructed by insertion of the BE3-blasticidin fragment into plasmid PB-CA (Addgene #20960). The BE3-blasticidin fragment was obtained by assembly of the BE3 fragment amplified from the pCMV-BE3 vector (Addgene #73021) and the blasticidin fragment amplified from lentiCas9-Blast (Addgene #52962). To establish a line of BE3-expressing cells, we co-transfected A375 cells with transposase (pPbase, Sanger Institute, UK) and the *piggyBac*-BE3-GFP plasmid at a 1:4 molar ratio using Lipofectamine 3000 (Invitrogen). At 48 h after transfection, the cells were cultured in medium containing 5 μg/mL of blasticidin for 14 days to enrich for BE3-expressing cells.

### sgRNA design

All sgRNAs were designed using an in-house program. We searched for all “GGs” and “AGs” as PAM sequences and considered sgRNA sequences in the form 5’-N_20_-PAM-3’ as candidates in the CDS region of every gene isoform. The candidates were examined to determine whether at least one “C” was in the activity window (4^th^ to 8^th^ position from the PAM-distal end of the protospacer) and that a “C” was in the CDS region. All possible mutations in the activity window were calculated, and the associated amino acid changes were classified as either silent, nonsense, or missense by reference to the codon frame and strand of the gene. All information regarding region and frame were obtained from a GTF file of hg19 downloaded from the UCSC Table Browser.

### Cell culture

A375 (ATCC) and HEK293T (ATCC) cells were maintained in RPMI medium (for A375 cells) or DMEM (for HEK293T cells) (Gibco) supplemented with 10% fetal bovine serum (Gibco) and penicillin/streptomycin (Gibco) in a humidified 5% CO_2_ incubator at 37°C. Solutions of vemurafenib (Selleckchem) were prepared by dissolving the colorless powder in DMSO (BioReagent grade; Sigma-Aldrich). DMSO was used as a vehicle control. To analyze the response to vemurafenib, cells were treated with either DMSO alone or 2 μM vemurafenib in DMSO.

### sgRNA-expressing lentivirus production and infection for testing of 420 sgRNA library and control experiments

Two oligos were ordered in the following form: (5’-CACCG-[20nt spacer sequence]-3’, 5’-AAAC-[reverse complementary sequence of 20nt spacer]-C-3’) for #159, #25, #354, non-targeting, and safe-targeting sgRNAs. Spacer sequences for non-targeting and safe-targeting sgRNAs are 5’-GTTATGCGACGTTCCGCATA-3’, 5’-GATCTTAGTGCCTTCACCAC-3’, respectively. We selected one unassigned guide (0None_none_UNA1_211552.253(*35*)) as a non-targeting sgRNA. For designing safe-targeting sgRNA, we first defined a safe region as an intersection of regions classified in inactive chromatin states (Quies, ReprPC or Het) from the Roadmap Epigenomics project database (http://www.roadmapepigenomics.org/). From this safe region, we picked a target sequence which has a PAM sequence and is close to the MAP2K1 gene. Oligos were then annealed and cloned into BsmBI-digested lenti-guide-puro or lentiCRISPR plasmids by ligation reaction. The lentivirus was then generated by co-transfecting the plasmid library, psPAX2 (Addgene #12260), and pMD2.G (Addgene #12259) into HEK293T cells using Lipofectamine 3000 (Invitrogen) according to the manufacturer’s protocol. A375 cells were transduced with the lentivirus at a multiplicity of infection (MOI) of 0.1–0.3 in each of three independent biological replicates. The cells were selected by culturing in medium containing either 1 μg/mL puromycin for 7 days, or DMSO or 2 μM vemurafenib for 14 days. All of these reactions were done independently on five sgRNAs.

### Dose response curve

After puromycin selection of cells individually transduced by #159, non-targeting, or safe-targeting sgRNA, cells were plated into 96-well plates (1 × 10^3^ cells/well with four biological replicates). After 16 h, media was replaced with media containing the indicated vemurafenib concentrations and refreshed every other day. After 3 days of treatment, cell viability was measured using thiazolyl blue tetrazolium bromide (Sigma-Aldrich). Cells were incubated with thiazolyl blue tetrazolium bromide solution at a final concentration of 0.45 mg/mL for 3.5 h. DMSO (GC grade; Sigma-Aldrich) was then added and the absorbance (570–630 nm) was measured on a microplate reader.

### Generation of sgRNA library plasmid, virus, and infection

Designed sgRNA sequences were synthesized using programmable microarrays (CustomArray, USA). Sequences are listed in **Data file S2**. Oligos were cleaved from the microarray and PCR amplified using chip_fwd and chip_rev. PCR cycling was performed as follows: 95°C for 3 min, 25 cycles of 98°C for 20 s, 56°C for 15 s, 72°C for 30 s, and 72°C for 1 min. To ensure high-yield coverage of the PCR products, eight repeats of the PCR reaction were conducted. The second PCR was performed using the chip_2nd_fwd and chip_2nd_rev primers. The PCR cycling conditions were 95°C for 3 min, followed by 6 cycles of 98°C for 20 s, 58°C for 15 s, 72°C for 30 s, and 1 cycle of 72°C for 1 min. The primers used in these steps are listed in Table S5. PCR products were purified using 2.0× (by volume) AMPure XP beads (Beckman Coulter). The purified amplicons were cloned into *Bsm*BI (NEB)-digested CROPseq-Guide-Puro plasmids (Addgene #86708) by Gibson assembly, as described previously.

To ensure high-yield coverage, four repeats of the Gibson reactions were performed. The products of the Gibson reactions were combined and electroporated into Endura cells (Lucigen) following the manufacturer’s protocol. A 1000-fold dilution of the full transformation was spread on carbenicillin (50 μg/mL) LB agar plates to determine the library coverage, and the remainder of the culture was incubated overnight in 200 ml of LB medium. By counting the number of colonies on the plate, >300× library coverage was ensured. The plasmid library was then extracted using an EndoFree Plasmid Maxi kit (Qiagen). The lentivirus was then generated by co-transfecting the plasmid library, psPAX2 (Addgene #12260), and pMD2.G (Addgene #12259) into HEK293T cells using Lipofectamine 3000 (Invitrogen) according to the manufacturer’s protocol.

A375 cells were transduced with the lentivirus at a multiplicity of infection of 0.1–0.3 in each of two independent biological replicates. The cells were selected by culturing in medium containing either 1 μg/mL of puromycin for 14 days or DMSO or 2 μM vemurafenib for 28 days.

### sgRNA-expressing lentivirus production and infection for validation of screening results

Two oligos were ordered in the following form: (5’-CACCG-[20nt spacer sequence]-3’, 5’-AAAC-[reverse complementary sequence of 20nt spacer]-C-3’) for #126, #56, and #217 sgRNAs. Oligos were then annealed and cloned into BsmBI-digested lenti-guide-puro or lentiCRISPR plasmids by ligation reaction. The lentivirus was then generated by co-transfecting the plasmid library, psPAX2 (Addgene #12260), and pMD2.G (Addgene #12259) into HEK293T cells using Lipofectamine 3000 (Invitrogen) according to the manufacturer’s protocol. A375 cells were transduced with the lentivirus at an MOI of 0.1-0.3 in each of two independent biological replicates. The cells were selected by culturing in medium containing either 1 μg/mL puromycin for 7 days, or DMSO or 2 μM vemurafenib for 14 days. All of these reactions were done independently on four sgRNAs.

### Preparation of sequencing library for 420 sgRNA targeting loci

Targeted sequencing for 420 sgRNA targeting regions was performed by Celemics Inc, Seoul, Korea. Briefly, 300 ng of gDNA was sheared into 300-bp fragments on average for adaptor-ligated library construction. Target sequences containing fragments were selected via hybrid capture using *MAP2K1*, *KRAS*, and *NRAS* targeting probes.

### Analysis of targeted capture sequencing data

From the NGS data, reads were aligned to hg19 using bwa v0.7.12-r1039 and then mapped reads were deduplicated with Picard version 1.128. Soft-clipped and indel-containing reads were excluded and base count for each ‘C’ position was calculated using pysam.

### Deep sequencing and analysis of sgRNA-integrated regions

To determine the frequency of each sgRNA, sgRNA sequences integrated into the genome were PCR amplified using the primers listed in Table S5. The resulting amplicons were adaptor-ligated using a SPARK kit (Enzymatics) and deep sequenced using NextSeq 500. Raw reads were quality trimmed via trimmomatic v0.33 using the following parameters: LEADING: 20, TRAILING: 20, SLIDINGWINDOW: 150:25, MINLEN: 36.

Reads containing each sgRNA spacer were counted, and these sgRNA read-counts were used as input for the MAGeCK software package to identify candidates by comparing DMSO-treated and vemurafenib-treated cells. The p-value of every sgRNA was calculated based on the negative binomial model of read counts. Log_2_(fold-change) was calculated as log 2 ratio of the normalized sgRNA count of vemurafenib-treated cells to that of DMSO-treated cells. Normalized sgRNA counts were calculated as reported previously(*3*).

### Deep sequencing of genomic DNA samples and C to T substitution efficiency

To validate whether the genome was edited by specific sgRNAs, the targeted region was amplified from the genome and sequenced. The primers used in these amplifications are listed in Table S5.

The PCR cycle conditions were as follows: 1 cycle at 95°C for 3 min, followed by 27 cycles of 98°C for 20 s, 56°C for 15 s, 72°C for 30 s, and 1 cycle of 72°C for 1 min. The resulting amplicons were adaptor-ligated and deep sequenced using NextSeq 500. If more than 50% of the bases that had a quality score lower than Q30, the sequenced reads were discarded, and base calls with a quality score below Q30 were converted to N. Ten-bp flanking sequences on both sides were used to identify the protospacer region for the targeted sgRNA. Protospacer and PAM sequence<u>-</u>containing reads that were not 23 bp in length were considered reads with indels. Base frequencies for each locus were calculated across the reads without indels. The plots are shown with the protein-coding strand, and if the protein-coding strand was the non-target strand of the sgRNA, G to A conversion efficiency was calculated (figs. S1B, S2D, S2E, S6J, S6K).

Substitution efficiencies were calculated using the following equation:

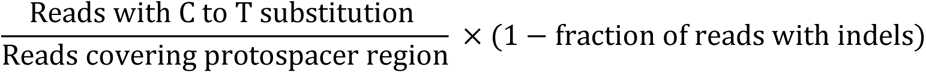

### scRNA-seq

A375 cells were harvested and divided into two pools on each sampling day. One pool was used for extraction of genomic DNA, and the other pool was methanol fixed for Drop-seq analysis. Methanol fixation was performed as described previously(*36*). Briefly, cells were centrifuged at 300*g* for 5 min and resuspended in 80% methanol (BioReagent grade; Sigma-Aldrich) and 20% PBS (Gibco). The resuspended cells were incubated on ice for at least 15 min and then stored at −80°C. On the day of the Drop-seq experiment, cells were recovered by centrifugation at 2000*g* for 5 min, washed once with PBS-0.01% BSA, and resuspended in PBS-0.01% BSA to a concentration of 100 cells/μl. Finally, Drop-seq was performed as described previously(*18*).

Droplets were collected into 50-ml conical tubes over a 15 min time period. After which, the droplets were broken, and the RNAs on beads were subjected to reverse transcription.

Aliquots of 2,000 beads were amplified in each tube using the following PCR steps: 95°C for 3 min, then four cycles of 98°C for 20 s, 65°C for 45 s, 72°C for 3 min, and 11 cycles of 98°C for 20 s, 67°C for 20 s, 72°C for 3 min, and 1 cycle of 72°C for 5 min. The amplified products were purified with 0.6× Ampure XP beads and fragmented, tagged, and amplified using a Nextera XT DNA Library Preparation kit (Illumina).

### Preprocessing of single-cell transcriptome data

The pipeline was designed based on Drop-seq tools (ver.1.12) and CROP-seq software. Raw data were converted to bam files via FastqToSam in Picard. Cell barcodes and UMI for each mRNA were obtained from Read1. mRNA sequences obtained from Read2 were modified by FilterBAM, TrimStartingSequence, and PolyATrimmer in Drop-seq tools. The modified bam files were then converted to fastq files and aligned to the reference comprised of hg19 and BE genes and guide RNA sequences using STAR aligner. The aligned data were sorted via SortSam and merged with tags via MergeBamAlignment in Picard. Exon information was annotated using TagReadWithGeneExon in Drop-seq tools, and bead synthesis errors within a hamming distance of <4 were corrected via DetectBeadSynthesisErrors in Drop-seq tools. For assignment of sgRNAs per each cell, UMI counts of each sgRNA in the same cell were determined by mapping the scRNA-seq reads to CROPseq-Guide-Puro plasmid chromosomes. If the most abundant sgRNA accounts for more than 90%, the cells were considered as uniquely assigned cell, and used for downstream transcriptome analysis. Finally, transcript data with more than 500 genes per cell were selected and converted to a digital gene expression matrix, and all matrices were merged and indexed by the cell barcode, condition, replicate, sgRNA, and gene. All program runs were managed using CROP-seq software(*13*).

### Estimation of MOI and detection rate

The initial MOI and detection rate were determined by the fitting method described previously(*12*). To reduce the selective effect, the data from initial cells for each replicate were only considered.

### Transcriptome analysis

Analyses were carried out based on the modules in Crop-seq software, Seurat(*37*), and Monocle 2(*38*).

#### Matrix modification for PCA and trajectory analysis

Merged matrices of digital gene expression were normalized per cell, pseudo-count 1 was added, multiplied by 10,000, and the matrix was finally log_2_ transformed. After normalization, ribosomal and mitochondrial genes and pseudo-references of sgRNAs and BE were filtered.

#### Dimensional reduction

First, PCA was performed using a PCA module in the sklearn package for global clustering. For characterization of D+28 samples, Seurat software was used. To follow the pipeline of Seurat, the original matrix before normalization was used and normalized again by Seurat’s internal functions. Briefly, cells from the D+28 samples treated with vemurafenib or DMSO with the appropriate fraction of mitochondrial genes (<0.065) were selected, and gene expression counts were normalized, multiplied by 10,000, and log_2_ transformed. Highly variable genes were selected using the FindVariableGenes function in the Seurat with the following parameters: mean.function = ExpMean, dispersion.function = LogVMR, x.low.cutoff = 0.05, x.high.cutoff = 3, y.cutoff = 1. The number of PCs was determined using PCElbowPlot and JackStaw. PCA was performed on selected variable genes using determined PCs. t-SNE was then applied.

#### Linear regression

Linear regression was performed for every gene using the statsmodel module in Python. Treatment, period (treated day), and their intersection were used as variables and fitted to the ordinary least square (OLS) model. In the case of treatment, a dummy variable was used as follows; 0 for the DMSO-treated condition, 1 for the vemurafenib-treated condition. Multiple testing corrections using the Benjamini and Hochberg method were applied with false discovery rate set to 0.05. Terms with corrected p-values of less than 0.05 were selected for every gene from the results. A heatmap was generated for visualization. Mean expression for guides were selected and plotted.

#### Euclidean distance

Euclidean distance was measured using the expression data of differential genes in the Vem(D+28) experiment. For distance between different sgRNAs targeting the same genomic position, we considered that sgRNAs target the same position when the target strands are identical and the protospacer regions overlap each other (**Data file S7**). Only distances from cells within the same replicate were calculated.

#### Gene set enrichment analysis

The Wilcoxon rank-sum test of the FindMarkers function in Seurat was used to assess differences in gene expression. Marker genes with low p-values (Benjamini -Hochberg(*39*) corrected p<0.05) for each cluster were obtained. GO and KEGG pathway enrichment analyses were carried out on the signature genes via clusterProfiler(*19*) and Enrichr(*20*). Multiple testing corrections using the Benjamini and Hochberg method with both p-value threshold and false discovery rate set to 0.05 were carried out.

#### Analysis of differentially expressed genes in DMSO-treated cells (DMSO(D+14) and DMSO(D+28)) with #176 aggregated from replicates 1 and 2, and #56 sgRNA from replicate 2

Two-sided Wilcoxon rank-sum test was performed to identify differentially expressed genes, and genes with low p-values (p < 0.05) were selected.

#### Trajectory Analysis

Trajectory analysis was performed by Monocle2. Expression matrix obtained from CROP-seq software were used as an input. Normalization and filtering were performed with parameters described in the “Filtering low-quality cells” section of Monocle2’s web document, and ordered genes were obtained using the differentialGeneTest module by setting the fullModelFormulaStr option. Finally, DDRTree-based dimensional reduction and ordering were performed.

### Specificity score

Specificity scores were defined based on the CFD model(*40*). For each sgRNA, the sum of CFD scores for all possible off-targets, of which the number of mismatches are less than 4, were added to 1 (score for the sgRNA itself). Then, a reciprocal of the value was used as a score as follows:

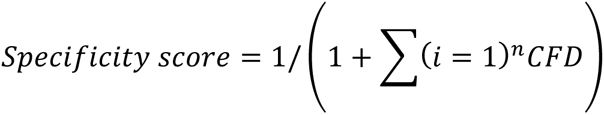

where *n* represents the number of possible off-target sites. The score would be 1 if the sgRNA is unique, and the score would be decreased when the sgRNA has off-targets. Of the possible off-targets in the coding region of the genome, protospacers which do not have a ‘C’ in the ‘activity window’ were excluded. Alternative PAMs except for AG were excluded due to low editing efficiency.

## Data Availability

Raw sequencing data from this study are deposited in Short Read Archive (SRA) under project number PRJNA530348. Data file S1 is deposited in Dropbox: https://www.dropbox.com/s/zqnu2qjyicllhdg/Data-File_S1.tsv?dl=0

## Acknowledgements

This work was supported by: (i) the Mid-career Researcher Program (NRF-2018R1A2A1A05079172), (ii) the Bio & Medical Technology Development Program (NRF-2016M3A9B6948494), (iii) the Bio & Medical Technology Development Program (NRF-2018M3A9H3024850), and (iv) Basic Science Research Program (NRF-2018R1A2B2001322) of the National Research Foundation of Korea, funded by the Ministry of Science, ICT & Planning, (v) Korea Health Technology R&D Project (HI18C2282) through the Korea Health Industry Development Institute (KHIDI), funded by the Ministry of Health & Welfare.

## Author contributions

S.J. and H.L. should be regarded as joint first authors. D.B. and J.H.L. conceived, designed, managed, and supervised the study. S.J. and H.L. designed and performed the experiments, analyzed the data, and wrote the manuscript. H.C. advised on Drop-seq device and provided PDMS chips for Drop-seq.

---

**Fig. S1.**
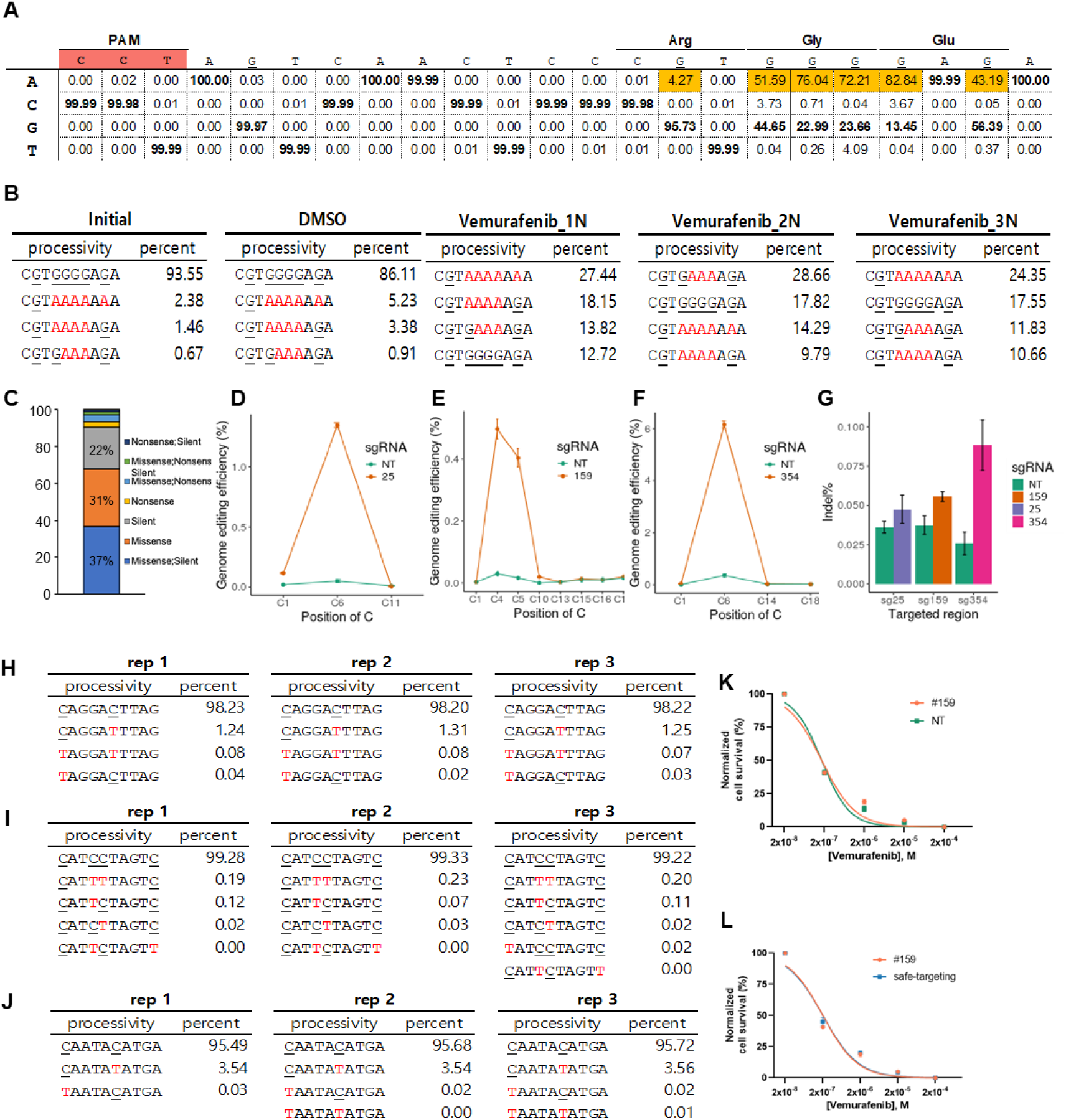
Conversions of C were observed within the ‘activity window,’ and processive conversions were observed. (**A**) Percentage of sequencing reads with the corresponding base is shown along with the sequence of the protospacer of the sgRNA. The plot is shown for the protein-coding strand, which is the non-target strand of the sgRNA; therefore, guanine was considered the target instead of cytosine. (**B**) Processive deamination was observed. The PAM-proximal 10 nt were examined for processivity. Convertible guanine is underlined, and converted adenine is shown in red font. To improve the confidence of the experimental enrichment for vemurafenib resistance, cells transduced by the virus in a single experiment were split into three sub-pools. Cell sub-pools were each treated with vemurafenib in triplicate. (**C**) Fraction of each introducible type of mutation using the 420-sgRNA library. (**D-F**) Efficiency of cytosine substitution according to position in the protospacer of the #25, #159, and #354 sgRNAs, respectively. Error bars represent the standard error of mean from three independent experiments. (**G**) Bar graph showing indel frequency in cells individually transduced by #25, #159, and #354 sgRNA. Error bars represent the standard error of mean from three independent experiments. Processive deamination was observed in singleplex experiments for (**H**) #25 sgRNA, (**I**) #159 sgRNA, and (**J**) #354 sgRNA. The PAM-distal 10 nt were examined for processivity. Convertible cytidine is underlined, and converted thymine is shown in red font. Vemurafenib dose response curves for (**K**) cells transduced with #159 sgRNA or non-targeting sgRNA (F_1,16_ = 0.1275, p = 0.7258, extra sum-of-squares F test between estimated logIC50, n = 2 replicates) and (**L**) cells transduced with #159 sgRNA or safe-targeting sgRNA (F_1,16_ = 0.4238, p = 0.5243, extra sum-of-squares F test between estimated logIC50, n = 2 replicates). The error bars represent the standard deviation from two independent experiments.

**Fig. S2.**
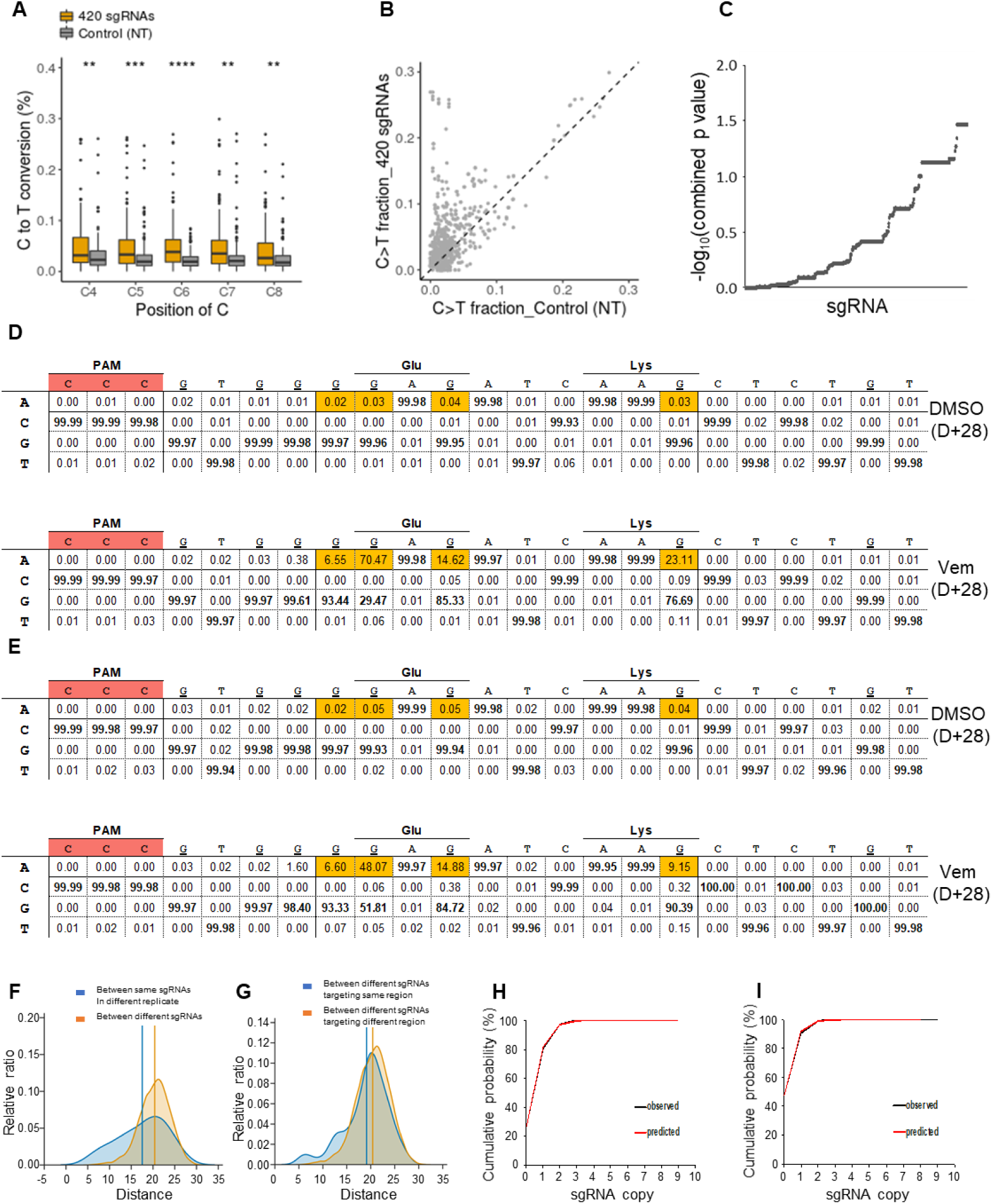
Introduction and functional screening of multiple mutations using population analysis. (**A-C**) Investigation of C to T conversion from 420 sgRNA-treated cells before drug selection. (**A**) Boxplot showing C to T conversion frequency within the ‘activity window’ for 420 sgRNAs. All p-values determined by two-sided Wilcoxon rank-sum test; ****p < 0.0001, ***p < 0.001, **p < 0.01, ns = not significant (p > 0.05). The center line corresponds to the median; the edges of the box correspond to the first and third quartiles; the whiskers represent 1.5× the interquartile range; and the black dots indicate outliers. (**B**) Plot of average C to T fraction of each “C” within the ‘activity window’. (**C**) Statistical significance of the C to T conversion ratio. Difference of C to T conversion between cells treated with 420 sgRNA and non-targeting sgRNA was tested with the one-sided Mann-Whitney U test and corrected using Benjamini-Hochberg method. P-values for each sgRNA were aggregated using Fisher’s method. Data for 420 sgRNA were produced using two independent replicate experiments, and data for non-targeting (NT) were produced using three independent replicates experiments. (**D**) Percentage of sequencing reads with the corresponding base is shown along with the sequence of the protospacer of the #176 sgRNA in replicate 1. The plot is shown for the protein-coding strand, which is the non-target strand of the sgRNA; therefore, guanine was considered the target instead of cytosine. (**E**) Percentage of sequencing reads with the corresponding base is shown along with the sequence of the protospacer of the #176 sgRNA in replicate 2. The plot is shown for the protein-coding strand, which is the non-target strand of the sgRNA; therefore, guanine was considered the target instead of cytosine. (**F-G**) Similarity between sgRNAs. (**F**) Distribution of distance between the same sgRNA in different replicates (blue) and different sgRNAs (red). (**G**) Distribution of distance between different sgRNAs targeting the same region (blue) and different sgRNAs targeting different regions (red). (**H-I**) Distribution of observed sgRNA copy number per cell (black) and the predicted value from the maximum likelihood estimate (red) for replicate 1 (**H**) and replicate 2 (**I**) samples.

**Fig. S3.**
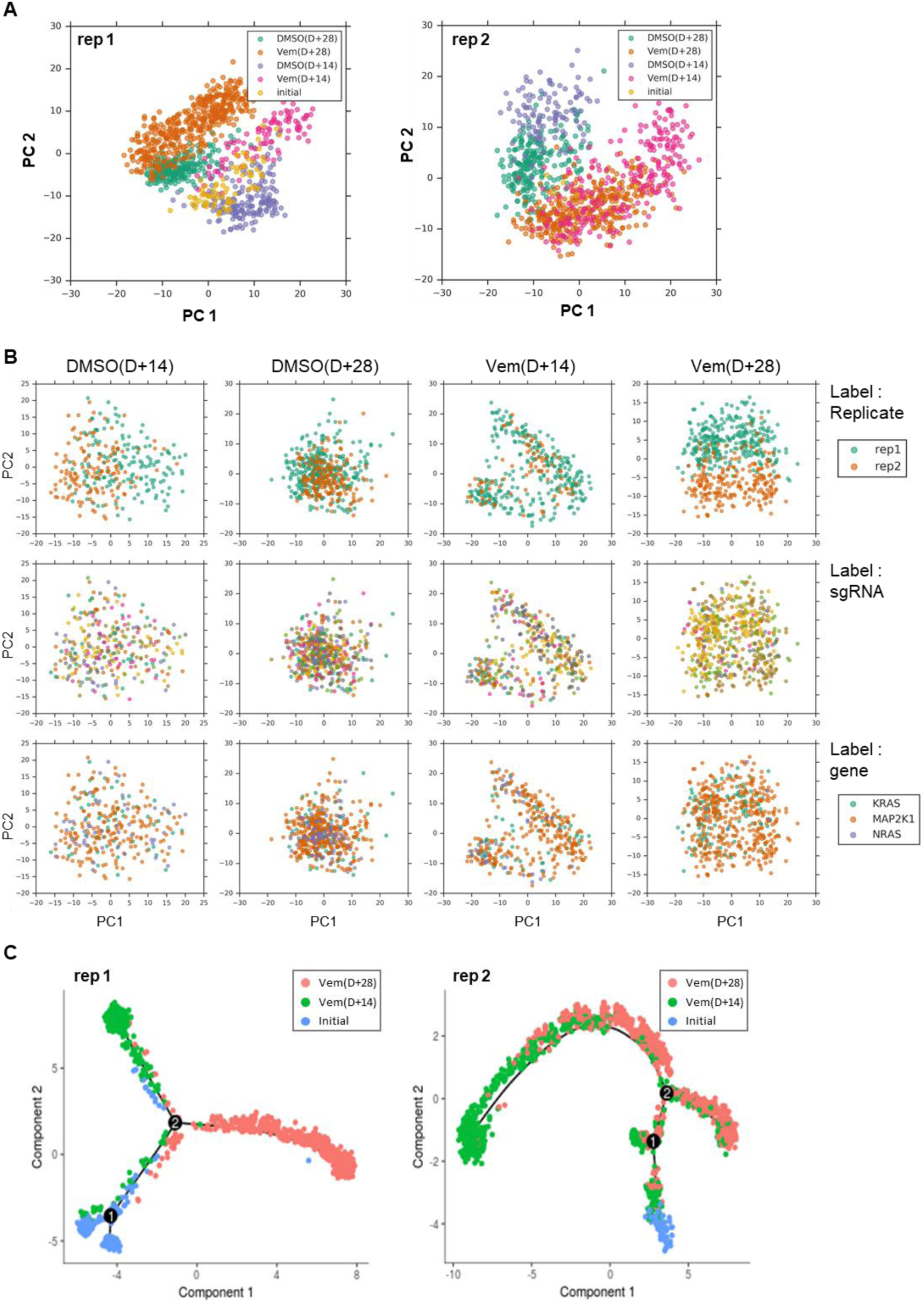
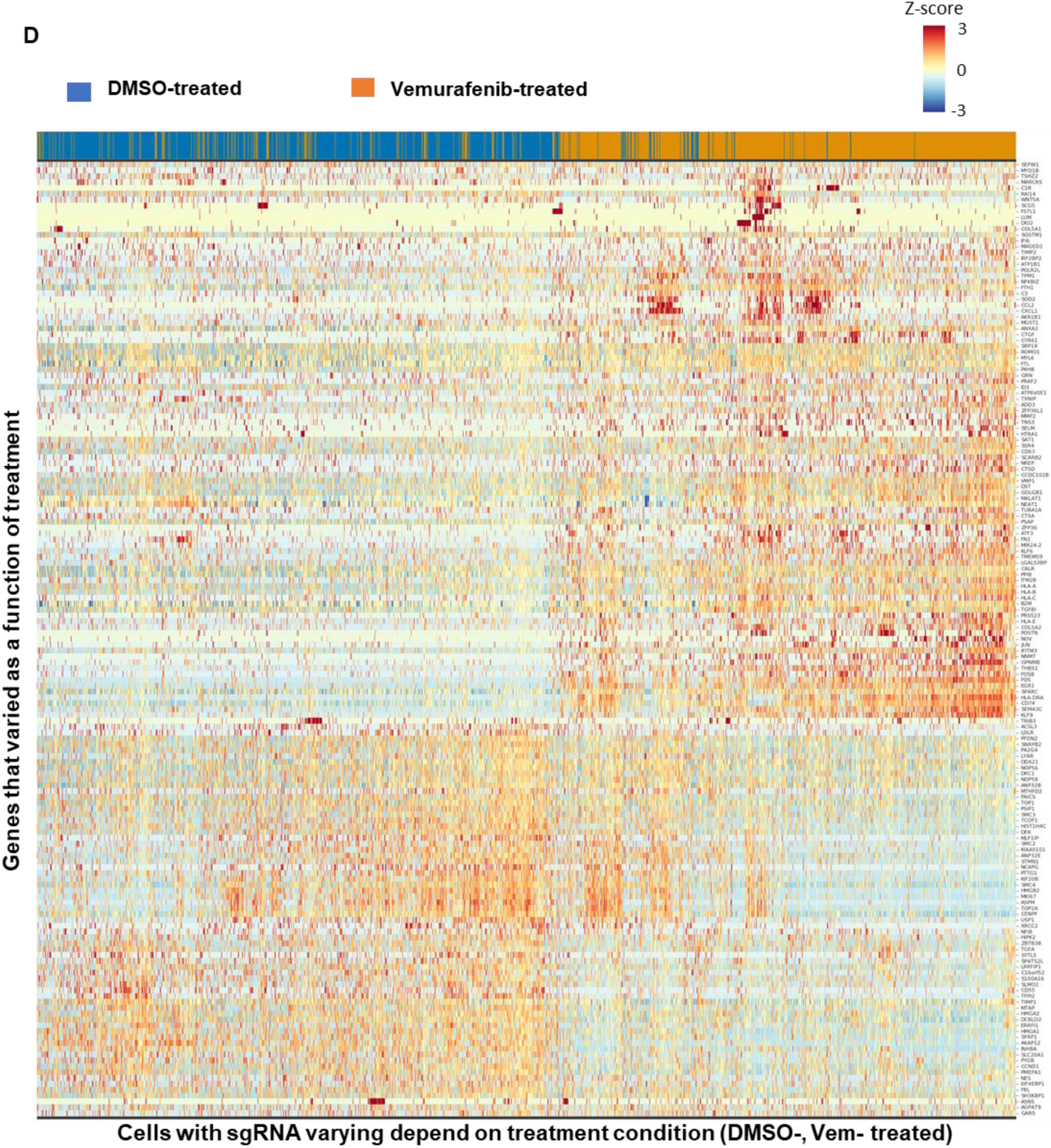
Global clustering across the different treatment periods. (**A**) PCA of transcriptomes for cells from the Initial, D+14, and D+28 data in replicate 1 (left) and replicate 2 (right). (**B**) PCA of transcriptomes for cells from DMSO(D+14), DMSO(D+28), Vem(D+14), and Vem(D+28). Cells were labeled according to replicate (top panel), sgRNA (middle panel), and gene (bottom panel). (**C**) Trajectory analysis using Monocle 2. The trajectories of Vem(D+14) and Vem(D+28) branched to different clusters in both replicates, and no progression according to treatment period was observed. (**D**) Average expression of genes, which were observed to vary with vemurafenib treatment, according to individual sgRNA. Each row represents genes and each column represents sgRNA. Color scale indicates the standard deviation of gene expression from the mean expression value, with red indicating high expression and blue indication low expression.

**Fig. S4.**
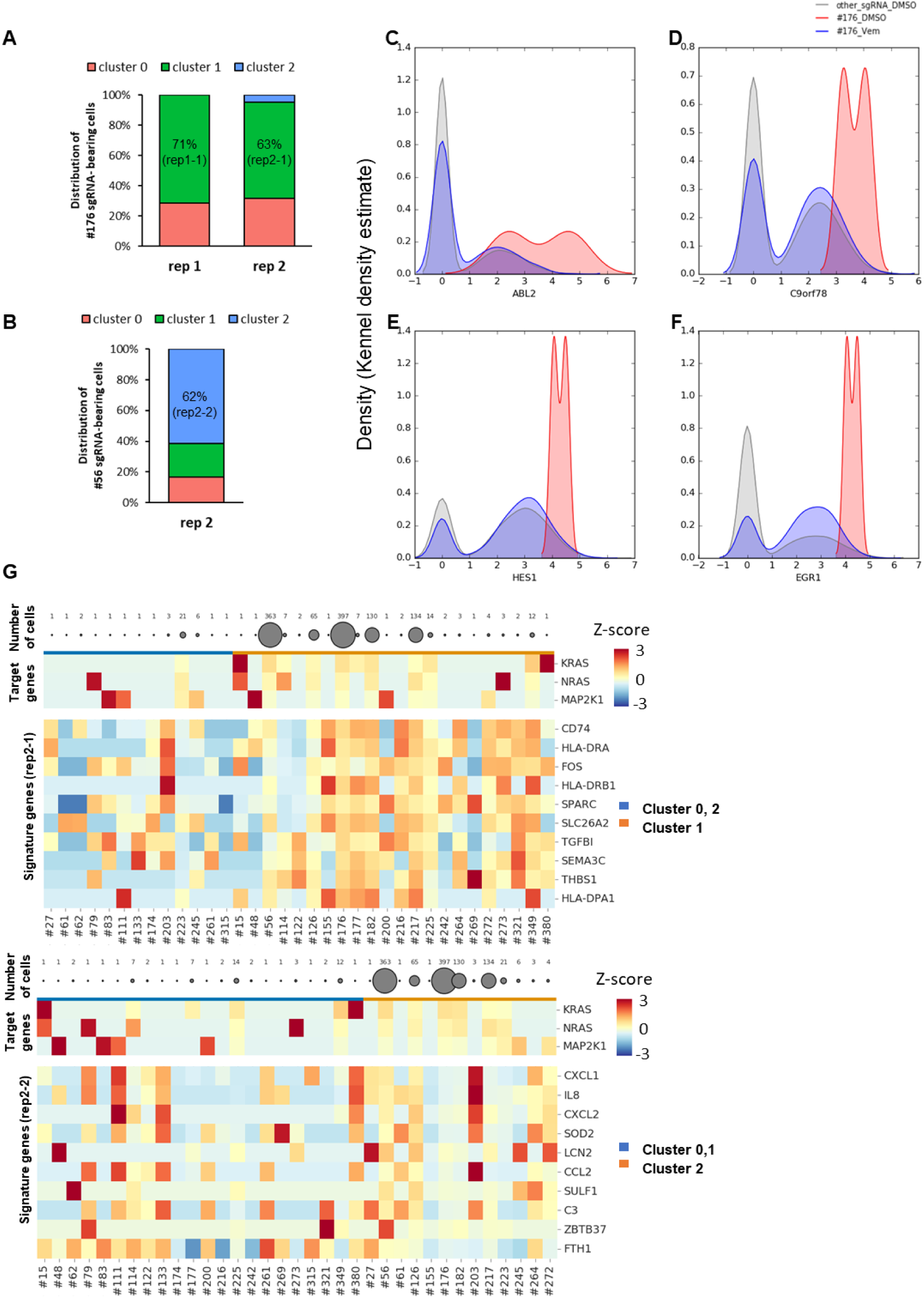
(**A**) Distribution of cells with the #176 sgRNA across clusters. Boxplot shows the fraction of #176 sgRNA located in each cluster. (**B**) Boxplot shows the fraction of #56 sgRNA located in each cluster. Expression of upregulated genes (**C**: *ABL2*, **D**: *C9orf78*, **E**: *HES1*, **F**: *EGR1*) in DMSO-treated cells with #176 sgRNA. (**G**) Profile of the transcription of signature genes in clusters composed primarily of the #176 sgRNA in replicate 2 (top) and profile of the transcription of signature genes in clusters composed primarily of #56 sgRNA in replicate 2 (bottom). Average expression of signature genes according to individual sgRNA. Each row represents one of the signature genes, and each column represents cells bearing the sgRNA. Color scale indicates the standard deviation of gene expression from the mean expression value, with red indicating high expression and blue indicating low expression.

**Fig. S5.**
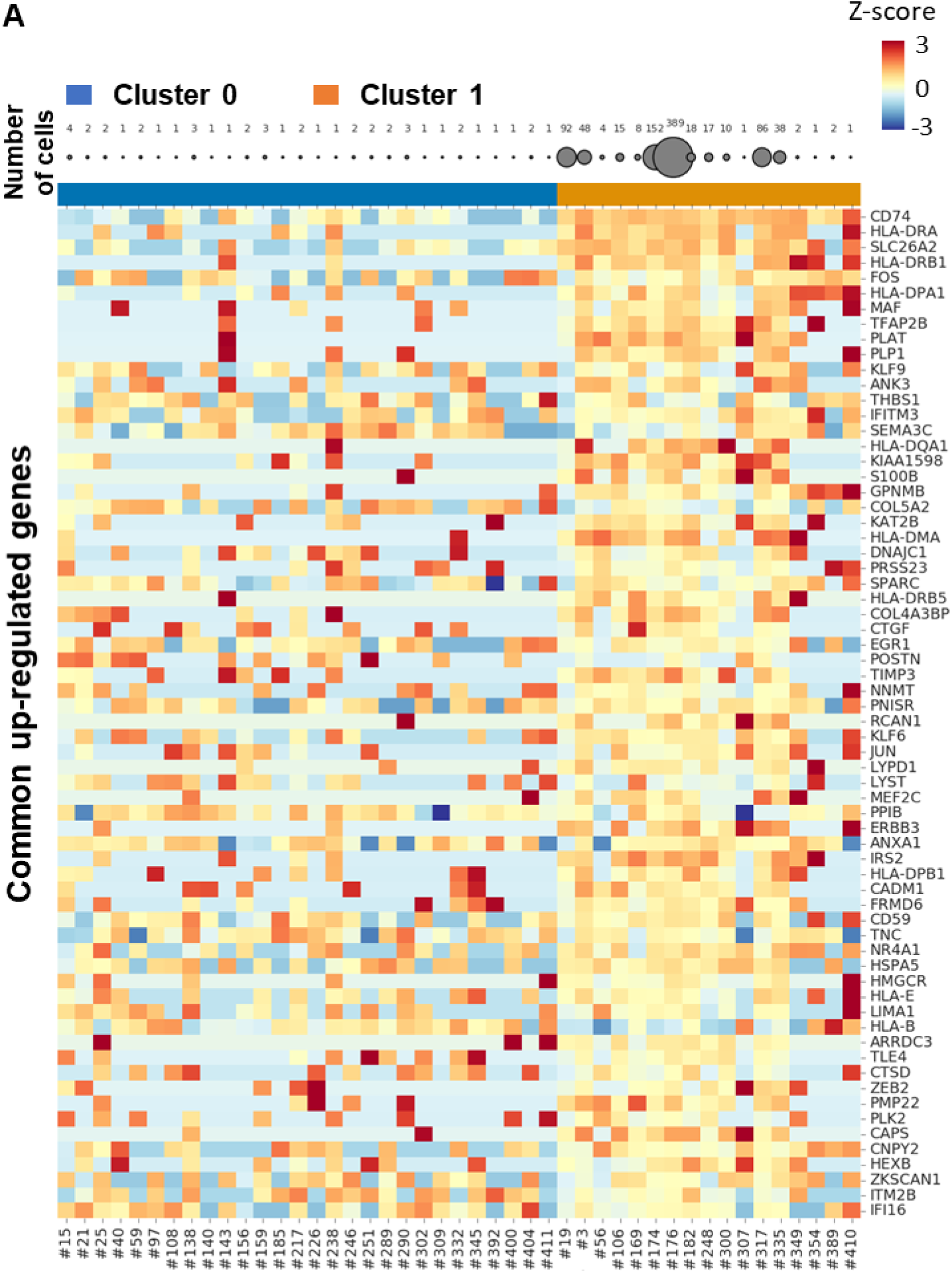
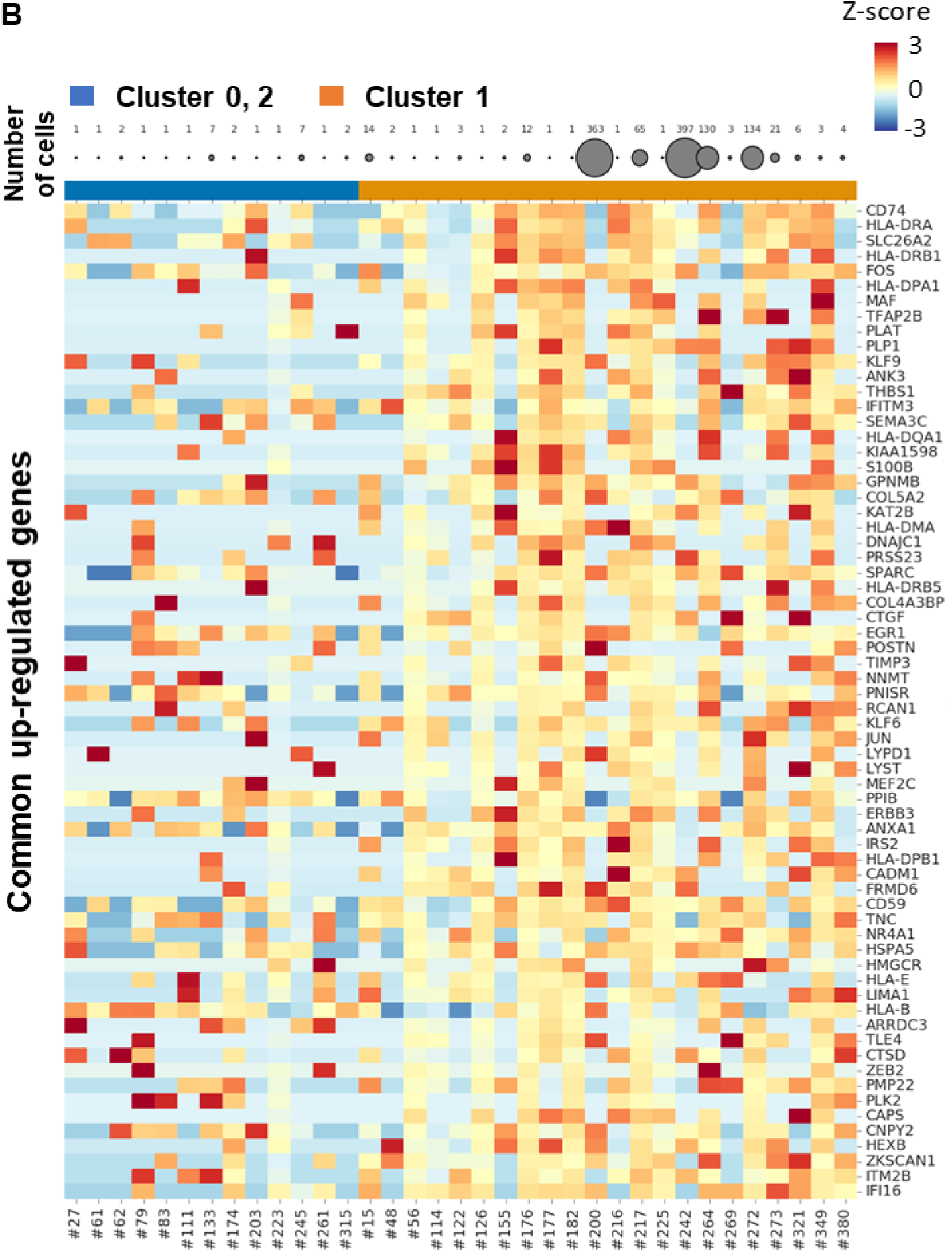
Average expression of signature genes according to individual sgRNA. Each row represents one of the common 66 up-regulated genes, and each column represents cells bearing the sgRNA. Cells from Vem(D+28) in other clusters or Vem(D+28) in rep1-1 (**A**) and rep2-1 (**B**) are arranged from left to right. Color scale indicates the standard deviation of gene expression from the mean expression value, with red indicating high expression and blue indicating low expression.

**Fig. S6.**
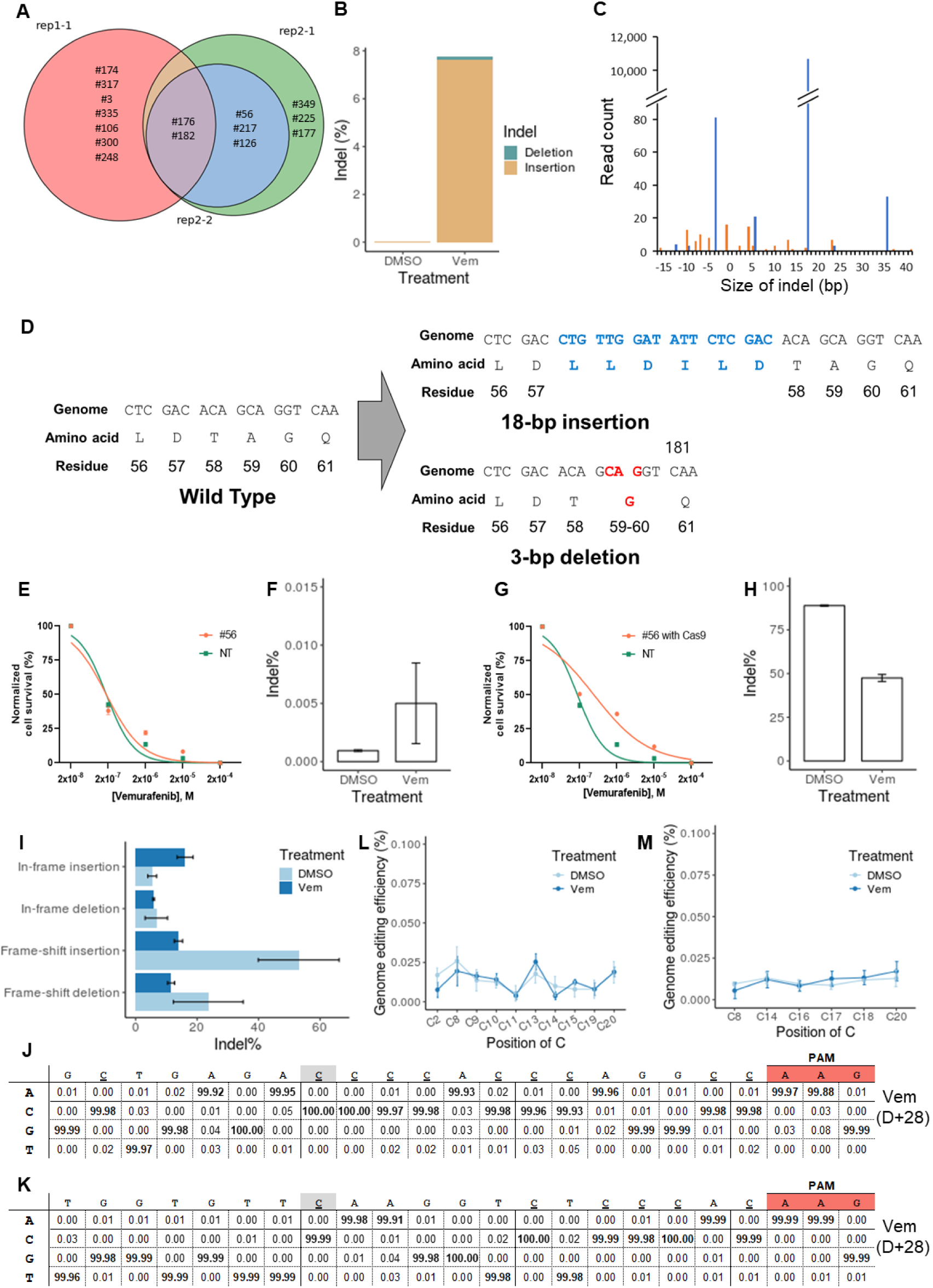
(**A**) Venn diagram of sgRNAs in each cluster that were contained in more than 2 cells. (**B-D**) Indel mutations found in the target region of the #56 sgRNA in replicate 2. (**B**) Bar graph showing indel frequency in cells treated with DMSO or vemurafenib for 28 days. (**C**) Distribution of read counts of the Indel mutations according to insertion size (positive value) and deletion (negative value). In-frame insertions or deletions are shown in blue, and frame-shift insertions or deletions are shown in orange. (**D**) Two major subtypes of indel mutations. Genome sequence of 18-bp insertion (right upper) is marked in blue, and that of the 3-bp deletion (right lower) is marked in red. Corresponding amino acid and position are shown beneath the genome sequence. (**E-I**) Singleplex experiments for #56 sgRNA. (**E**) Vemurafenib dose response curves (F_1,26_ = 0.03237, p = 0.8586, extra sum-of-squares F test between estimated logIC50, n = 3 replicates) and (**F**) bar graph showing indel frequency for cells expressing BE3 and #56 sgRNA. (**G**) Vemurafenib dose response curves (F_1,26_ = 13.94, p = 0.0009, extra sum-of-squares F test between estimated logIC50, n = 3 replicates) and (**H**) bar graph showing indel frequency for cells expressing Cas9 and #56 sgRNA. (**I**) Bar graph showing frame-shift or in-frame indel frequency. p-value for significance of enrichment was determined by one-sided Wilcoxon rank-sum test, *p < 0.05. Error bars represent the standard error of mean from three independent experiments. (**J-K**) Conversion of C by the #126 and #217 sgRNAs was not observed within the “activity window” in replicate 2. Percentage of sequencing reads with the corresponding base is shown along with the sequence of the protospacer of the #126 (**J**) and #217 (**K**) sgRNAs. Cytosines within the activity window are shown in grey. (**L-M**) Singleplex experiments for #126 and #217 sgRNAs. (**L**) Efficiency of cytosine substitution according to position in the protospacer of the #126 sgRNA. (**M**) Efficiency of cytosine substitution according to position in the protospacer of the #217 sgRNA. All experiments were performed in three independent replicates. Error bars represent the standard error of mean from three independent experiments.

**Table S1.** Detailed information regarding the study data. Table shows the characteristics of scRNA-seq data for each step of the study.

|  | D0 |  | D+14 |  |  |  | D+28 |  |  |  |
| --- | --- | --- | --- | --- | --- | --- | --- | --- | --- | --- |
|  | Initial |  | DMSO |  | Vemurafenib |  | DMSO |  | Vemurafenib |  |
|  | rep1 | rep2 | rep1 | rep2 | rep1 | rep2 | rep1 | rep2 | rep1 | rep2 |
| Sequenced reads (*10<sup>6</sup>) | 45.01 | 1.47 | 76.29 | 130.59 | 150.65 | 97.54 | 62.37 | 88.50 | 71.47 | 95.67 |
| Uniquely mapped reads % | 59.99 | 66.73 | 54.54 | 42.46 | 26.26 | 54.12 | 58.18 | 52.14 | 50.62 | 55.56 |
| Multiple mapping reads % | 15.57 | 15.75 | 14.10 | 12.08 | 7.96 | 13.90 | 13.86 | 12.79 | 13.27 | 14.44 |
| Cells with >500 genes | 1,606 | 107 | 1,003 | 1,560 | 940 | 2,341 | 627 | 1,543 | 1,531 | 1,960 |
| Mean unique reads per cell | 2,260 | 1,614 | 3,534 | 3,811 | 2,289 | 3,114 | 4,959 | 5,125 | 4,032 | 4,048 |
| Mean genes per cell | 1,166 | 1,066 | 1,651 | 1,613 | 1,249 | 1,494 | 1,950 | 2,069 | 1,772 | 1,823 |
| gRNA assigned cells with >500 genes (unique) | 849 | 83 | 619 | 847 | 516 | 1,252 | 324 | 755 | 971 | 1,212 |
| Unique gRNA assignment % (cells with >500 genes) | 52.86 | 77.57 | 61.71 | 54.29 | 54.89 | 53.48 | 51.67 | 48.93 | 63.42 | 61.84 |

**Table S2.**
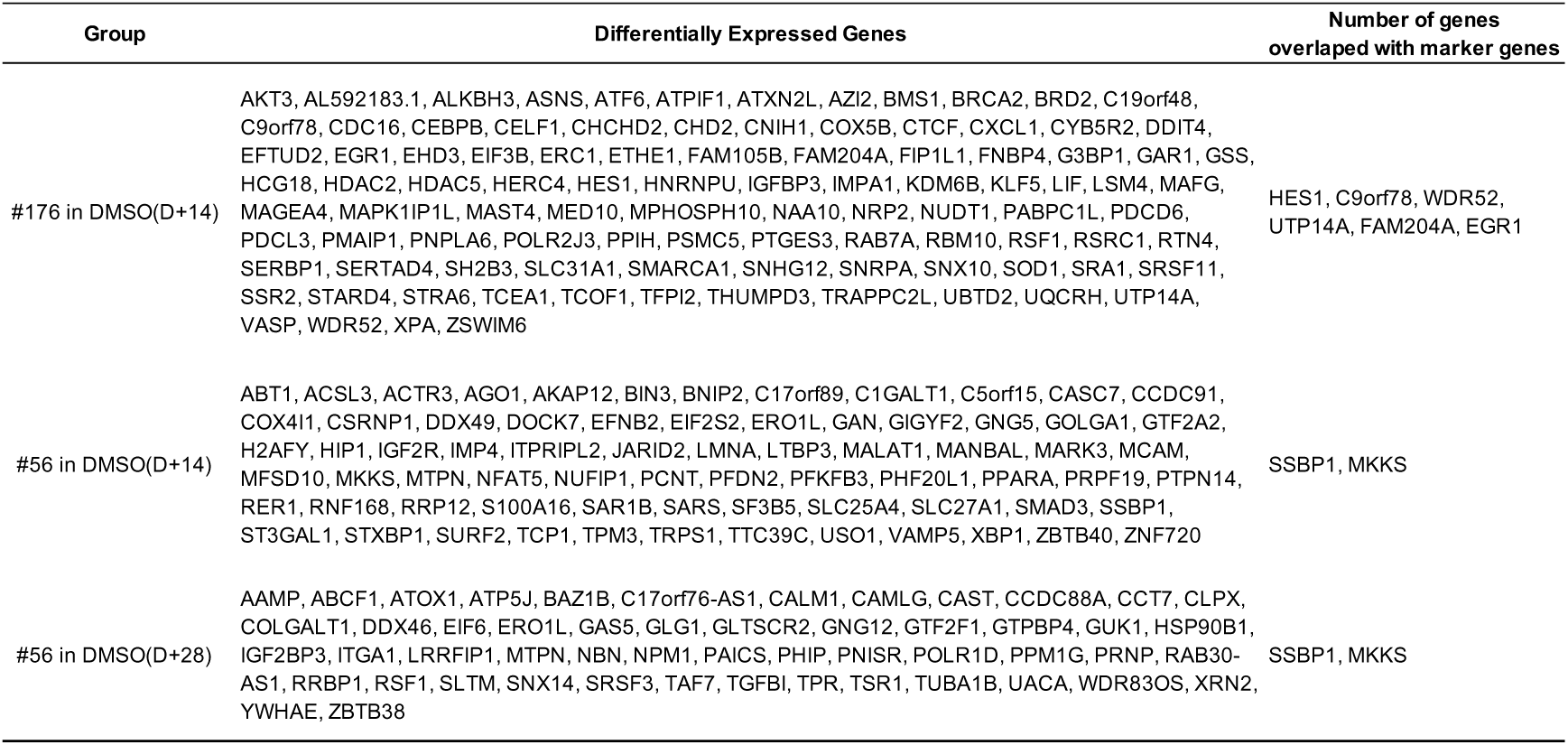
Genes that varied under DMSO-treated conditions in cells with #176 and #56 sgRNAs.

**Table S3.** sgRNAs in the rep1-1, rep2-1, and rep2-2 clusters. Corresponding counts are shown to the right of each sgRNA.

| Cluster | sgRNA | Count | Cluster | sgRNA | Count | Cluster | sgRNA | Count |
| --- | --- | --- | --- | --- | --- | --- | --- | --- |
| rep1-1 | #176 | 105 | rep2-1 | #176 | 154 | rep2-2 | #56 | 135 |
|  | #174 | 29 |  | #56 | 48 |  | #126 | 19 |
|  | #317 | 26 |  | #182 | 47 |  | #176 | 12 |
|  | #3 | 13 |  | #217 | 46 |  | #182 | 8 |
|  | #335 | 8 |  | #126 | 11 |  | #217 | 4 |
|  | #182 | 6 |  | #349 | 6 |  | #272 | 2 |
|  | #106 | 4 |  | #225 | 6 |  | #61 | 1 |
|  | #300 | 3 |  | #177 | 3 |  | #203 | 1 |
|  | #248 | 3 |  | #216 | 2 |  | #297 | 1 |
|  | #169 | 2 |  | #272 | 2 |  | #223 | 1 |
|  | #19 | 2 |  | #200 | 1 |  | #348 | 1 |
|  | #389 | 1 |  | #242 | 1 |  | #368 | 1 |
|  | #410 | 1 |  | #155 | 1 |  | #155 | 1 |
|  | #347 | 1 |  | #15 | 1 |  | #27 | 1 |
|  | #307 | 1 |  | #269 | 1 |  | #245 | 1 |
|  | #56 | 1 |  | #380 | 1 |  | #264 | 1 |
|  | #349 | 1 |  | #114 | 1 |  |  |  |
|  | #354 | 1 |  | #273 | 1 |  |  |  |
|  |  |  |  | #48 | 1 |  |  |  |
|  |  |  |  | #321 | 1 |  |  |  |
|  |  |  |  | #264 | 1 |  |  |  |
|  |  |  |  | #122 | 1 |  |  |  |

**Table S4.**
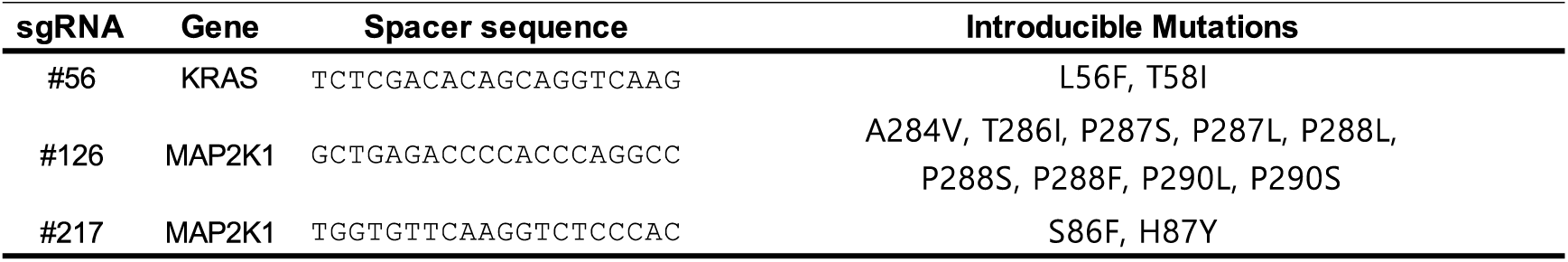
Cognate mutations of candidate genes.

**Table S5.**
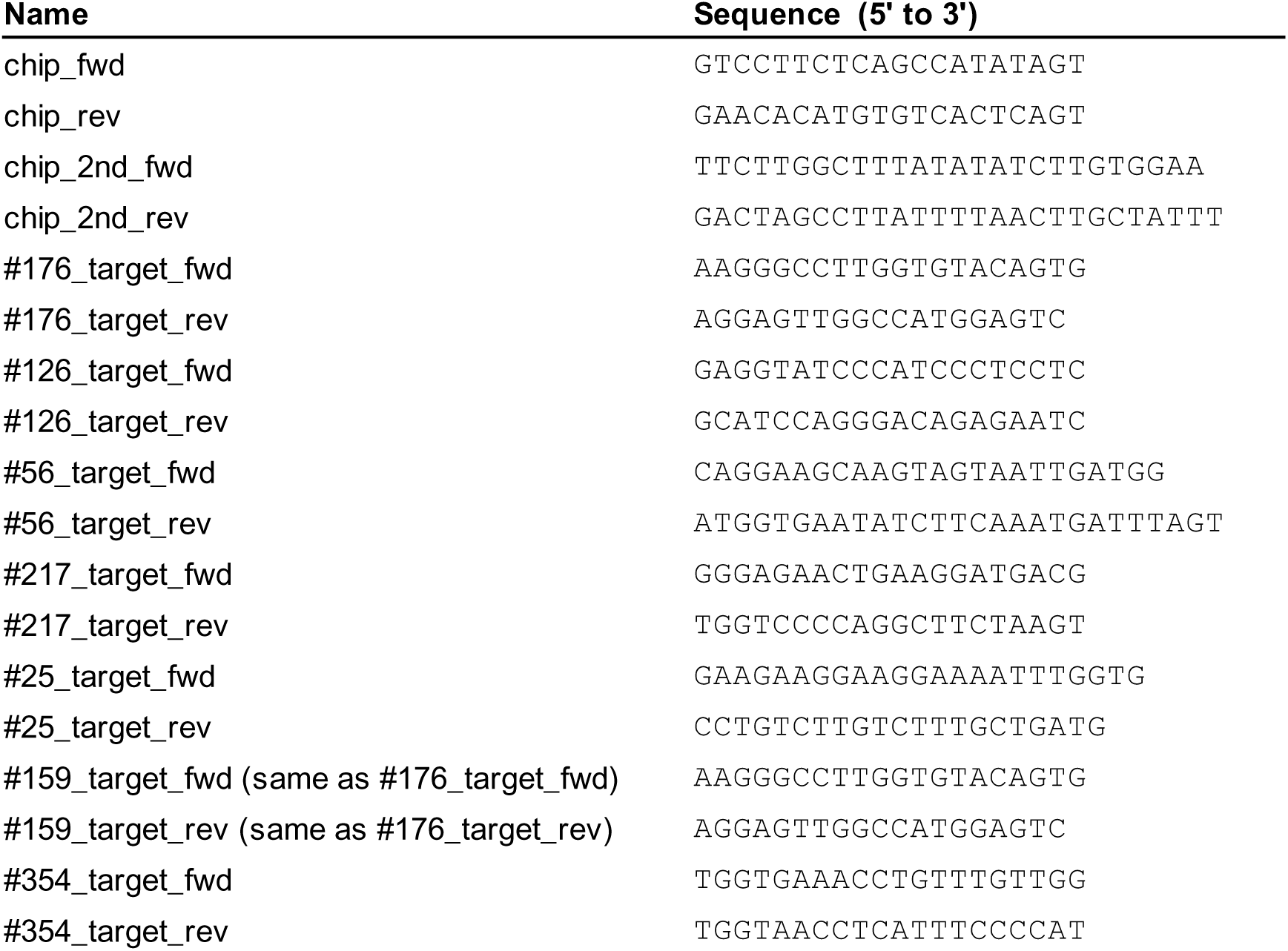
Primers used in this study.

